# Nej1^XLF^ interacts with Sae2^CTIP^ to inhibit Dna2 mediated resection at DNA double strand break

**DOI:** 10.1101/2021.04.10.439283

**Authors:** Aditya Mojumdar, Nancy Adam, Jennifer A. Cobb

## Abstract

The two major pathways of DNA double strand break (DSB) repair, non-homologous end-joining (NHEJ) and homologous recombination (HR), are highly conserved from yeast to mammals. The regulation of 5’ DNA resection controls repair pathway choice and influences repair outcomes. Nej1 was first identified as a canonical NHEJ factor involved in stimulating the ligation of broken DNA ends and more recently, it was shown to be important for DNA end-bridging and inhibiting 5’ resection mediated by Dna2-Sgs1. Nej1 interacts with Sae2 and this impacts DSB repair in three ways. First, Nej1 inhibits MRX-Sae2 interactions and Sae2 localization to a DSB. Second, Nej1 inhibits Sae2-dependent recruitment of Dna2 in the absence of Sgs1. Third, *NEJ1* and *SAE2* showed an epistatic relationship for DNA end-bridging, an event that restrains the broken ends and reduces the frequency of genomic deletions from developing at the DSB. Deletion of *NEJ1* suppressed the synthetic lethality of *sae2*Δ *sgs1*Δ and was dependent on the nuclease activity of Dna2. These Nej1 functions promote end-joining DSB repair, but could also be relevant for controlling resection initiation during HR repair.

**Highlights:**

- Nej1 physically interacts with Sae2 and inhibits end-resection at a DSB.
- Nej1 inhibits Sae2 interactions with the MRX complex.
- Nej1 inhibits Sae2-dependent recruitment of Dna2 to a DSB.
- NEJ1 and SAE2 are epistatic for DNA end-bridging.
- Deletion of *NEJ1* suppresses the synthetic lethality of *sae2*Δ *sgs1*Δ, which is dependent on Dna2 nuclease activity.

## 1. Introduction

DNA double-strand breaks (DSBs) can be repaired by two central pathways, non-homologous end joining (NHEJ) and homologous recombination (HR). NHEJ mediates the direct ligation of DNA ends without the requirement for end processing, whereas HR requires 5’ end resection. Both 5’ resection and end-bridging are important for repair pathway choice and down-stream DNA processing. Once resection initiates at a DSB, repair by canonical NHEJ is no longer an option. A network of proteins, involving the core NHEJ factor Nej1, has evolved to regulate this key step [1-9].

yKu70/80 (Ku) and Mre11-Rad50-Xrs2 (MRX) are the first complexes that localize to a DSB and both are important for recruiting Nej1 [1-4]. Cells lacking *NEJ1* are as defective in end-joining repair as *ku70*Δ and *dnl4*Δ [3, 5, 6]. Nej1 contributes to Ku stability, which protects the DNA ends from nucleolytic degradation, and promotes Lif1-Dnl4 mediated ligation [2, 3, 7, 8]. Nej1 also functions in collaboration with MRX to bridge DNA ends at the DSB. The structural features of the MRX complex are critical for end-bridging and deletion of *NEJ1* result in end-bridging defects that are additive with *rad50* mutants [4, 10-13]. While Nej1 and MRX both contribute to DNA end-bridging, Nej1 functions antagonistically to MRX by inhibiting 5’ DNA resection. Currently, few mechanistic details exist for how Nej1 inhibits 5’ resection, although earlier work showed that Nej1 inhibits Dna2 interactions with Sgs1 and Mre11 [4]. Its role in repair pathway choice certainly involves more than reinforcing Ku-dependent DNA end protection for NHEJ. In fact, a certain level of Ku must be maintained at DSBs in *nej1*Δ mutants as increased 5’ resection was Dna2 dependent and Exo1 independent [4, 9].

5’ DNA resection occurs through a two-step process [14]. First, Sae2, the yeast homologue of human CtIP, activates Mre11 endonuclease to initiate DNA resection, which also promotes Ku dissociation from the DNA ends [15, 16]. Second, long-range resection follows and is mediated by two functionally redundant 5’ to 3’ nucleases, Dna2, in complex with Sgs1, and Exo1 [16, 17]. Mre11 endonuclease activity is less critical for 5’ resection than the physical presence of the MRX complex at DSBs. Exo1 and Dna2-Sgs1 can serve as compensatory back-ups to initiate resection, however both long-range nucleases require MRX for localization [9, 17, 18]. Exo1 has a very high affinity for DNA ends that are not protected by Ku and it can initiate resection in *mre11* nuclease dead (nd) mutants only when *KU70* is deleted, but not when *NEJ1* was deleted [19, 20].

The regulation of Dna2 in resection is more complex than that of Exo1 and understanding its entire function at DSBs has been challenging because *DNA2* is an essential gene involved in Okazaki fragment processing, therefore it cannot be deleted [21-25]. Earlier work showed that the lethality of *dna2*Δ can be suppressed by disruption of *PIF1* helicase, and that the frequency of 5’ resection decreased at a DSB in *dna2*Δ *pif1*-m2 mutants [19, 26]. In the absence of Mre11 nuclease activity, resection initiates primarily through Dna2, independently of *KU* status [27-29]. Moreover, using nuclease deficient *dna2*-1 (P504→S), Dna2 and Mre11 showed functional redundancy for processing the ends of DSBs after radiation treatment [30]. Most work describing Dna2 at DSBs has been performed in surrogate, by deleting *SGS1* [16, 31]. However, those studies cannot explain the greater IR and UV sensitivity of *dna2*-1 *sgs1*Δ mutants compared to single mutant counterparts [32], and would not be able to identify any potential function(s) for Dna2 at DSBs independently of Sgs1. Another pathway of Dna2 recruitment to DSB is through human CtIP [33]. Sae2 was recently shown to stimulate the nuclease and helicase activity of Dna2-Sgs1 *in vitro*, although this needs to be demonstrated *in vivo* at the break site [34, 35]. Moreover, Sae2 also has a role in DNA end-bridging at DSBs [36], a function conserved in humans and with Ctp1 in fission yeast [37, 38]. Owing to their known functions, and the roles of Sae2 and Nej1 in promoting and inhibiting resection respectively, an investigation of the Nej1-Sae2 relationship at DSB is warranted.

In the current work, we demonstrate that Nej1 functions in opposition to Dna2 and Sae2 in DNA end processing at DSBs. The deletion of *NEJ1* led to increased 5’ resection and increased recovery of Dna2 and Sae2 at the break. We show that Sae2 is a key factor in Dna2 recruitment to DSBs. Nej1 binds with Sae2, inhibiting its physical association with each component of the MRX complex and with Dna2. Aside from regulating 5’ resection, Nej1-Sae2 interactions might be functionally important for restricting the mobility DNA ends at the break site because cells harboring *NEJ1* and *SAE2* deletions showed epistatic end-bridging defects. Lastly, we show that deletion of *NEJ1* can suppress the synthetic lethality of *sae2*Δ *sgs1*Δ through a mechanism dependent on the nuclease activity of Dna2.

## 3. Results

### 3.1. Nej1 Inhibits Sae2 recovery at a DSB

The recruitment of Sae2 and Nej1 to a DSB depends on the MRX complex [4, 28]. Sae2 initiates DNA end-resection by activating Mre11 endonuclease [16]. By contrast, Nej1 interacts with the C-terminus of Mre11 and inhibits resection [4,9], however the potential impact of Nej1 on Sae2 has not been determined. We performed chromatin immuno-precipitation (ChIP) on Sae2 with primers located 0.6 kb from the HO-DSB (Fig 1A). Consistent with previous work, Sae2 decreased to background levels in *mre11*Δ mutants (Fig 1B). By contrast, Sae2 recovery increased ∼2-fold in *nej1*Δ mutants from 40 mins to 3 hours after HO induction (Figs 1B,C). This was not an indirect consequence of disrupting NHEJ repair in general because Sae2 did not increase in cells where *KU70* or *DNL4* was deleted (Fig 1B). We also assessed the importance of Sae2 in Nej1 localization given that Ku70 recovery increases in *sae2*Δ mutants (Fig S1A, [19]). No change in Nej1 recovery was seen in *sae2*Δ mutants whereas its recovery in *mre11*Δ was reduced to background (Fig.1 D).

**Fig 1.**
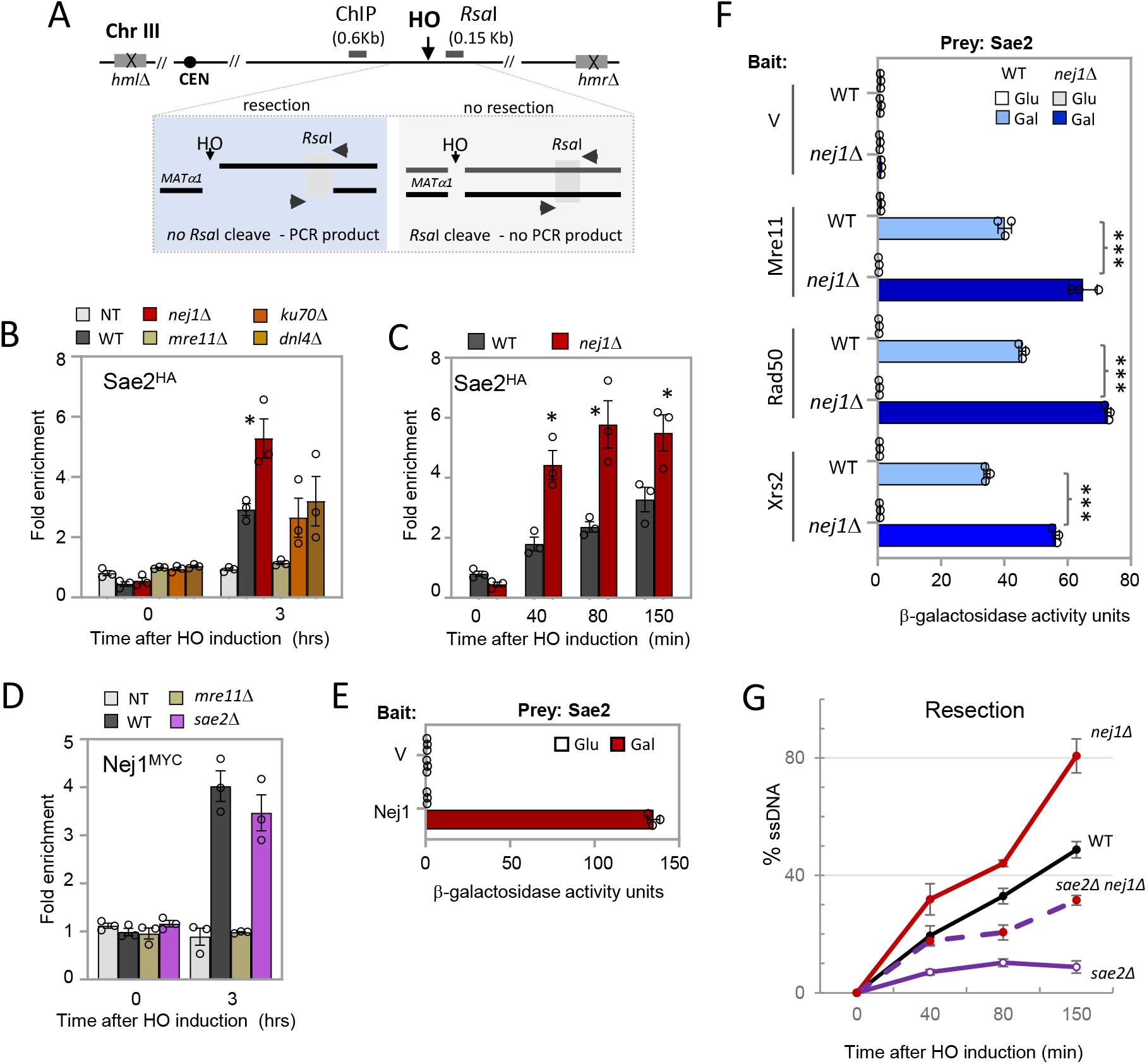
Sae2 recruitment at DSB is inhibited by Nej1. **(A)** Schematic representation of regions around the HO cut site on chromosome III. The ChIP probe used in this study is 0.6kb from the DSB. The RsaI sites used in the qPCR resection assays, 0.15kb from the DSB, are also indicated. **(B)** Enrichment of Sae2^HA^ at DSB, at 0 and 3 hours, in wild type (JC-5116), *nej1*Δ (JC-5124), *mre11*Δ (JC-5122), *ku70*Δ (JC-5948), *dnl4*Δ (JC-5946) and no tag control (JC-727). The fold enrichment represents normalization over the SMC2 locus. **(C)** Enrichment of Sae2^HA^ at 0.6kb from DSB, at 0 min (no DSB induction), 40, 80 and 150 mins after DSB induction in wild type (JC-5116) and *nej1*Δ (JC-5124). **(D)** Enrichment of Nej1^Myc^ at DSB, at 0 and 3 hours, in wild type (JC-1687), *mre11*Δ (JC-3677), *sae2*Δ (JC-5118) and no tag control (JC-727). **(E and F)** Y2H analysis of Sae2 fused to HA-AD, and Mre11, Rad50, Xrs2, and Nej1 fused to LexA-DBD was performed in wild type cells (JC-1280) and in isogenic cells with *nej1*Δ (JC-4556) using a quantitative β-galactosidase assay. **(G)** 5’ DNA Resection 0.15kb away from the HO DSB using a qPCR-based approach described in the methods section. Frequency of resection is plotted as % ssDNA at 0, 40, 80 and 150 mins. post DSB induction in cycling cells in wild type (JC-727), *nej1*Δ (JC-1342), *sae2*Δ (JC-5673) and *sae2*Δ *nej1*Δ (JC-5675). For all the experiments -error bars represent the standard error of three replicates. Significance was determined using 1-tailed, unpaired Student’s t test. All strains compared are marked (P<0.05*; P<0.01**; P<0.001***) and are compared to WT, unless specified with a line.

To determine whether there was a physical interaction between Nej1 and Sae2 we next performed yeast two-hybrid (Y2H) as previously described [4,8]. This approach was used because Nej1 has a short half-life making coimmunoprecipitation (co-IP) methods difficult [4, 5, 8, 39, 40]. Sae2 was expressed as hemagglutinin (HA) -tagged prey and Nej1 was expressed as LexA-tagged bait [4, 8, 41]. Sae2 showed robust binding with Nej1 upon galactose-induction (Figs 1E and S1B). We also performed Y2H between Sae2 and each component of the MRX complex. Consistent with previous reports [42], Sae2 physically interacted with the MRX complex (light blue bars, Fig 1F), and all interactions increased in *nej1*Δ mutants (dark blue bars, Fig 1F and S1B). Western blots showed that constructs expressed similarly in WT and *nej1*Δ backgrounds after galactose-induction (Fig S1C, D). The binding of Nej1 and Sae2 could inhibit the interactions between Sae2 with Mre11, Rad50, and Xrs2 when they were expressed as LexA-tagged bait. Thus, when a DSB occurs, Nej1 could inhibit Sae2 recruitment in two ways. First, by directly binding to Sae2, and secondly by interacting with MRX, which we previously mapped to the C-terminus (423-692 aa) of Mre11 [4].

Given that Sae2 promotes resection whereas Nej1 inhibits it, we next measured 5’ resection directly at the DSB using a quantitative PCR-based approach developed by others and previously used by us [4, 9, 36, 43]. It relies on an *Rsa*I cut site located 0.15 kb from the DSB (Fig 1A). If resection has proceeded past this site, then single-stranded DNA is produced, and the region can be amplified by PCR using primers that flank the restriction site. Deletion of *SAE2* reversed the elevated rate of 5’ resection in *nej1*Δ mutants (Fig 1G). The increased rate of resection in *nej1*Δ was dependent on a pathway involving Sae2 as *nej1*Δ *sae2*Δ double mutants showed reduced resection below wild type but above *sae2*Δ single mutants. Taken together with the ChIP and Y2H (Fig.1B-F), these data suggest that Nej1 inhibits Sae2-mediated resection at a DSB.

### 3.2. Nej1 regulates resection and HR by inhibiting Dna2 and Sae2

When Mre11 nuclease is not activated, as in *sae2*Δ mutants, resection initiates primarily from the activity of Dna2-Sgs1. These findings, together with our previous work showing Nej1 inhibits Dna2-Sgs1 [4], prompted us to determine Dna2 recovery in nuclease-dead *mre11*-3 mutants. The MRX complex is recovered similarly in *MRE11*+ and *mre11*-3 mutants, which is important as the other nucleases, Exo1 and Dna2 require it for their recruitment [44, 45]. Dna2 recovery in *mre11*-3 was indistinguishable from wild type (Fig. 2A), and was ∼2-fold above wild type in *mre11*-3 *nej1*Δ and *nej1*Δ mutants (Fig.2A, [4]). Consistent with increased Dna2 recovery, *mre11*-3 *nej1*Δ mutants also showed increased resection (Fig.2B). Dna2-Sgs1 functions as a complex in long-range resection and because *DNA2* is essential, *sgs1*Δ has been used as a surrogate for assessing Dna2 functionality at DSBs. Indeed, increased resection in *mre11*-3 *nej1*Δ and nej1Δ mutants was Sgs1 dependent as resection proceeded in *mre11*-3 *nej1*Δ *sgs1*Δ and *nej1*Δ *sgs1*Δ mutants similar to wild type (Fig.2B, D). Surprisingly, Dna2 recovery remained high in the triple mutants (Fig. 2A). In fact, Dna2 increased in all genetic combinations where *SGS1* was deleted if *NEJ1* was deleted also (Fig. 2A, C). In *sgs1*Δ single mutants, the recovery level of Dna2 decreased however, levels remained well above the non-tagged (NT) control (Fig. 2C). Taken together these results suggest an Sgs1-independent pathway for Dna2 recruitment that is also inhibited by Nej1. While resection is indeed more defective in *sgs1*Δ *exo1*Δ compared to *sgs1*Δ (Figs. 2D and S2A), the deletion of *EXO1* did not alter Dna2 recruitment to the DSB (Fig. 2C and S2B) nor did it reverse the hyper-resection phenotype of *nej1*Δ mutants, even when Mre11 activity was abrogated in *mre11*-3 *nej1*Δ mutants (Fig. S2C, D). Furthermore, the recovery of Exo1 did not change in *nej1*Δ like it did in *ku70*Δ mutants when end protection is lost (Fig. S2E). In all, our data suggests the presence of an alternate Sgs1-independent pathway for Dna2 recruitment, which is inhibited by Nej1, and does not depend on Exo1. Given the interactions between Nej1 and Sae2, we next measured Sae2 recovery in these various mutants. While Sae2 recruitment was abrogated in *mre11*Δ (Fig 1B), its localization increased in *mre11*-3 mutants (Fig. 2E), which is consistent with earlier work [28]. Conversely, in *sgs1*Δ and *exo1*Δ mutants, DSB-associated Sae2 remained indistinguishable from wild type (Fig. 2F). Sae2 recovery increased ∼5-fold in *mre11*-3 *nej1*Δ, which was higher than in either single mutant, and this increase was sustained in *mre11*-3 *nej1*Δ *sgs1*Δ triple mutants (Fig 2E). In all, our data suggest that the Sgs1-independent pathway for Dna2 recruitment involves Sae2. Sae2 recovery was not impacted by deletion of *SGS1* (Figs. 2E, F), increased levels of both Dna2 and Sae2 correlate with increased resection in *mre11*-3 *nej1*Δ *sgs1*Δ, compared to *mre11*-3 *sgs1*Δ double mutants which have decreased resection and comparatively less Dna2 and Sae2 recovery.

**Fig 2.**
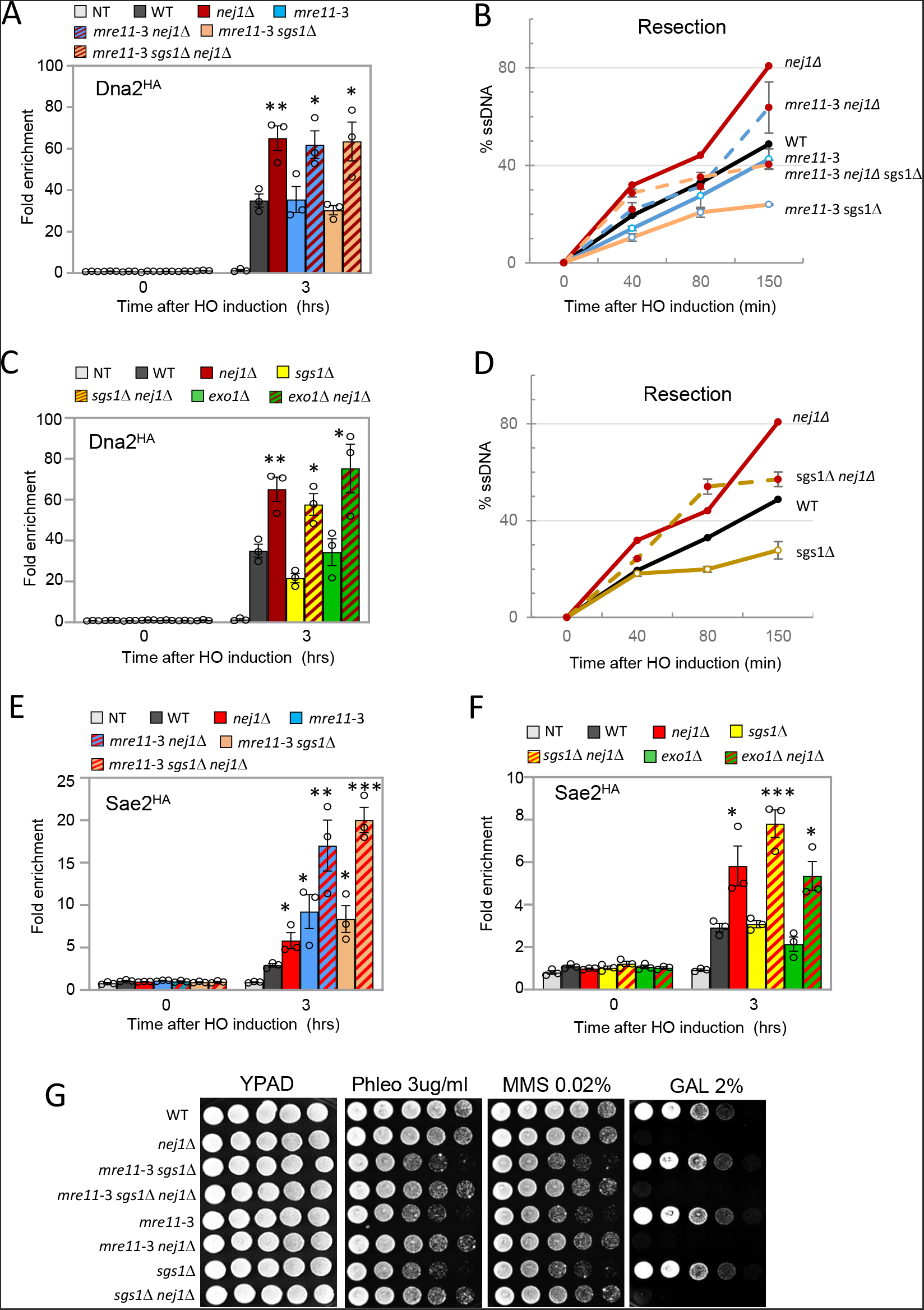
Nej1 regulates resection and HR by inhibiting Dna2 and Sae2. **(A and C**) Enrichment of Dna2^HA^ at 0.6kb from DSB, at 0 and 3 hours after DSB induction in wild type (JC-4117), *nej1*Δ (JC-4118), *mre11*-3 (JC-5594), *mre11*-3 *nej1*Δ (JC-5596), *mre11*-3 *sgs1*Δ (JC-5621), *mre11*-3 *sgs1*Δ *nej1*Δ (JC-5623), *sgs1*Δ (JC-5624), *sgs1*Δ *nej1*Δ (JC-5627), *exo1*Δ (JC-5626), *exo1*Δ *nej1*Δ (JC-5666) and no tag control (JC-727) was determined. The fold enrichment is normalized to recovery at the SMC2 locus. **(B and D)** 5’ DNA Resection 0.15kb away from the HO DSB using a qPCR-based approach described in the methods section. Frequency of resection is plotted as % ssDNA at 0, 40, 80 and 150 mins. post DSB induction in cycling cells in wild type (JC-727), *nej1*Δ (JC-1342), *mre11*-3 (JC-5372), *mre11*-3 *nej1*Δ (JC-5369), *mre11*-3 *sgs1*Δ (JC-5405), *mre11*-3 *sgs1*Δ *nej1*Δ (JC-5667), *sgs1*Δ (JC-3757) and *sgs1*Δ *nej1*Δ (JC-3759). **(E and F)** Enrichment of Sae2^HA^ at DSB, at 0 and 3 hours, in wild type (JC-5116), *nej1*Δ (JC-5124), *mre11*-3 (JC-5119), *mre11*-3 *nej1*Δ (JC-5702), *mre11*-3 *sgs1*Δ (JC-5704), *mre11*-3 *sgs1*Δ *nej1*Δ (JC-5706), *sgs1*Δ (JC-5684), *sgs1*Δ *nej1*Δ (JC-5685), *exo1*Δ (JC-5688), *exo1*Δ *nej1*Δ (JC-5686) and no tag control (JC-727) was determined. The fold enrichment is normalized to recovery at the SMC2 locus. **(G)** Five-fold serial dilutions of the strains in (B and D) were spotted on YPAD, Phleomycin 3.0 ug/ml, MMS 0.02% and Galactose 2%.

Furthermore recent reports show that Sae2 can stimulate the nuclease activity of Dna2 [34, 35]. Survival on Phleomycin and MMS, two agents that cause DNA double strand breaks DSBs, was greater in *mre11*-3 *nej1*Δ *sgs1*Δ than *mre11*-3 *sgs1*Δ mutants, suggesting that increased Dna2 and/or resection led to increased DSB repair, when it can proceed via HR. These strains also contained one HO cut site and survival after DSB induction by galactose, at a location where HR repair is precluded, showed the opposite trend in survival and demonstrated the requirement of Nej1 in end-joining repair (Fig 2G).

### 3.3. Nej1 interactions with Sae2 regulate Dna2 recruitment and end-bridging

Resection was lower in *sae2*Δ *nej1*Δ and *sae2*Δ compared to *mre11*-3 *nej1*Δ and *mre11*-3 respectively (Figs 1G and 2B), which is consistent with Sae2 having functions in DSB repair beyond Mre11 nuclease activation [28]. We wanted to determine whether these additional functions included Sae2-dependent recruitment of Dna2 to the DSB, and whether this was inhibited by Nej1 given higher levels of Dna2 and Sae2 were recovered in *nej1*Δ and *mre11*-3 *nej1*Δ mutants independently of SGS1 status. Increased Dna2 recovery in *mre11*-3 *nej1*Δ double mutants was Sae2 dependent as the enrichment level in *mre11*-3 *nej1*Δ *sae2*Δ was similar to wildtype (Fig, 3A). Moreover, deletion of *SAE2* also reversed the elevated resection in *mre11*-3 *nej1*Δ mutants (Fig, 3B). The recovery of Dna2 decreased in *sae2*Δ and *mre11-3 sae2*Δ mutant cells, even more than in *sgs1*Δ and *mre11*-3 *sgs1*Δ (Fig 3A). In drop assays *mre11*-3 *nej1*Δ *sae2*Δ mutants showed no greater resistance to Phleomycin or MMS compared to *mre11*-3 *sae2*Δ (Fig. 3C). These data suggest that increased Dna2 levels at DSBs, along with increased resection by *NEJ1* deletion, was important for reversing the Phleomycin and MMS sensitivity of *mre11*-3 *sgs1*Δ (Fig. 2G)

**Fig 3.**
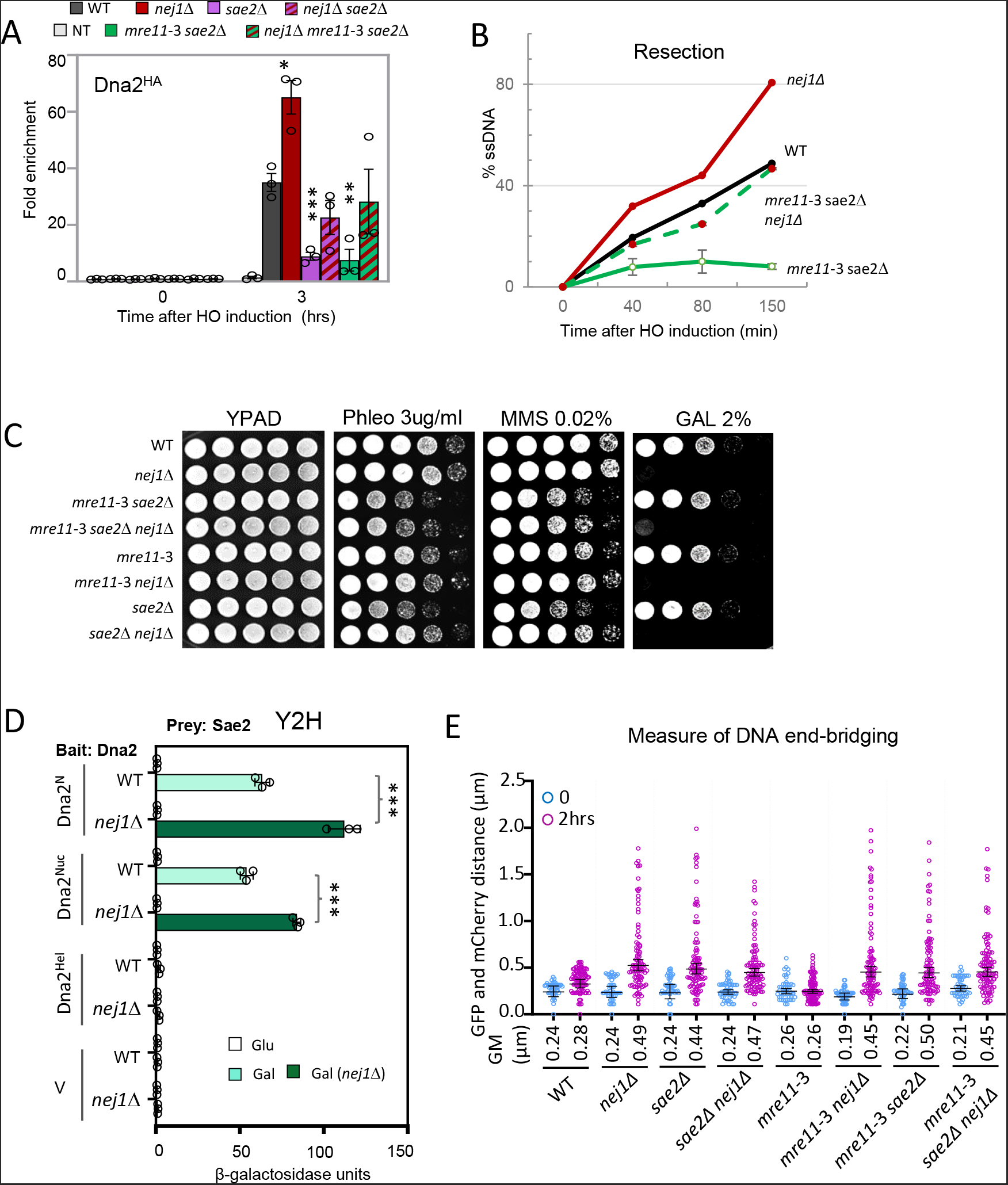
Sae2-dependent recruitment of Dna2 is inhibited by Nej1. **(A)** Enrichment of Dna2^HA^ at 0.6kb from DSB 0 hour (no DSB induction) and 3 hours after DSB induction in wild type (JC-4117), *nej1*Δ (JC-4118), *sae2*Δ (JC-5562), *sae2*Δ *nej1*Δ (JC-5597), *sae2*Δ *mre11*-3 (JC-5598), *sae2*Δ *mre11*-3 *nej1*Δ (JC-5593) and no tag control (JC-727) was determined. The fold enrichment is normalized to recovery at the SMC2 locus. **(B)** 5’ DNA Resection 0.15kb away from the HO DSB using a qPCR-based approach described in the methods section. Frequency of resection is plotted as % ssDNA at 0, 40, 80 and 150 mins. post DSB induction in cycling cells in wild type (JC-727), *nej1*Δ (JC-1342), *sae2*Δ *mre11*-3 (JC-5501) and *sae2*Δ *mre11*-3 *nej1*Δ (JC-5500). **(C)** Five-fold serial dilutions of the cells in wild type (JC-727), *nej1*Δ (JC-1342), *sae2*Δ *mre11*-3 (JC-5501) and *sae2*Δ *mre11*-3 *nej1*Δ (JC-5500), *mre11*-3 (JC-5372), *mre11*-3 *nej1*Δ (JC-5369), *sae2*Δ (JC-5673) and *sae2*Δ *nej1*Δ (JC-5675) were spotted on YPAD, Phleomycin 3.0 ug/ml, MMS 0.02% and Galactose 2%. **(D)** Y2H analysis of Sae2 fused to HA-AD and domains of Dna2, (Dna2-N terminal, Dna2-Nuclease and Dna2-Helicase domains) fused to LexA-DBD was performed in wild type cells (JC-1280) and in isogenic cells with *nej1*Δ (JC-4556) using a quantitative β-galactosidase assay. **(E)** Scatter plot showing the tethering of DSB ends, at 0 and 2 hours, as measured by the distance between the GFP and mCherry foci in wild type (JC-4066), *nej1*Δ (JC-4364), *sae2*Δ (JC-5524), *sae2*Δ *nej1*Δ (JC-5525), *mre11*-3 (JC-5529), *mre11*-3 *nej1*Δ (JC-5526), *sae2*Δ *mre11*-3 (JC-5530) and *sae2*Δ *mre11*-3 *nej1*Δ (JC-5531). The Geometric mean (GM) distance for each sample is specified under the respective sample data plot. Significance was determined using Kruskal-Wallis and Dunn’s multiple comparison test. For all the experiments -error bars represent the standard error of three replicates. Significance was determined using 1-tailed, unpaired Student’s t test. All strains compared are marked (P<0.05*; P<0.01**; P<0.001***) unless specified with a line, strains are compared to WT.

These data suggest that Sae2 functions with Dna2 to promote resection. Therefore, we next determined whether Sae2 and Dna2 physically interacted. Yeast-two-hybrid (Y2H) was performed as previously described with prey vector HA-tagged Sae2 expressed together with the following bait vectors: LexA-tagged Dna2 domains - 1-450 aa (N-term), 451-900 aa (nuclease) and 901-1522 aa (helicase) as previously described (Fig. S3A) [4, 41]. Sae2 interacted with the N-terminal regulatory region and nuclease domains of Dna2 (light green bars). Similar to Sae2-MRX, Sae2 interactions with Dna2^N^ and Dna2^Nuc^ increased in *nej1*Δ mutants (dark green bars; Figs 3D and S3B). Of note, deletion of *NEJ1* did not increase general binding between all proteins expressed from 2-hybrid vectors as Mre11-Rad50 interactions were unaltered in *nej1*Δ cells and all constructs were similarly expressed in WT and *nej1*Δ backgrounds (Fig S3C-E). Taken together with previous work [4], these data suggest that Nej1 functions as a general inhibitor of interactions between nucleases and their binding partners. Nej1 inhibits both Sae2-MRX and Sae2-Dna2 interactions in addition to Dna2-Sgs1 and Dna2-Mre11 interactions [4].

Nej1 is essential for end-joining, therefore growth on galactose was markedly reduced in all mutant combinations containing *nej1*Δ as seen in drop assays on 2% galactose (Figs. 2G and 3C) and in more quantitative cell survival measurements (Table 1). Survival on continuous galactose inversely correlates with 5’ DNA resection and in all cells, wild type and mutants. Cell survival under these conditions is very low because only cells that have acquired mutations that prevent recutting will survive (Table 1). The mating type of survivors is very informative as it provides insight about *in vivo* DNA processing events, revealing genomic alterations that develop at the HO-DSB located within MATα1 and adjacent to MATα2 (Fig S4A). Expression of these genes regulates the mating type by activating α-type genes and inhibiting a-type genes. Extensive resection leading to large deletions (>700 bp) produces ‘a-like’ survivors because both α1 and α2 are disrupted [9]. Consistent with previous reports, large deletions developed in *nej1*Δ (Table 1 and Fig S4A). The frequency of survivors that developed large deletions in *nej1*Δ mutants was partly reduced by further deleting SGS1 or SAE2, which also correlated with their ability to reverse the elevated rate of resection in *nej1*Δ mutants. Large deletion also decreased, but to a lesser extent when *nej1*Δ was combined with *mre11*-3, but not in combination with deletion of *EXO1*, as resection remained elevated in *nej1*Δ *exo1*Δ (Fig 2SC and Table 1). Similar trends were also seen in *mre11*-3 *nej1*Δ *sgs1*Δ and *mre11*-3 *nej1*Δ *sae2*Δ triple mutant survivors which showed a further decrease in the frequency of large deletions compared to *mre11*-3 *nej1*Δ (Table 1 – S4A).

**Table 1.**
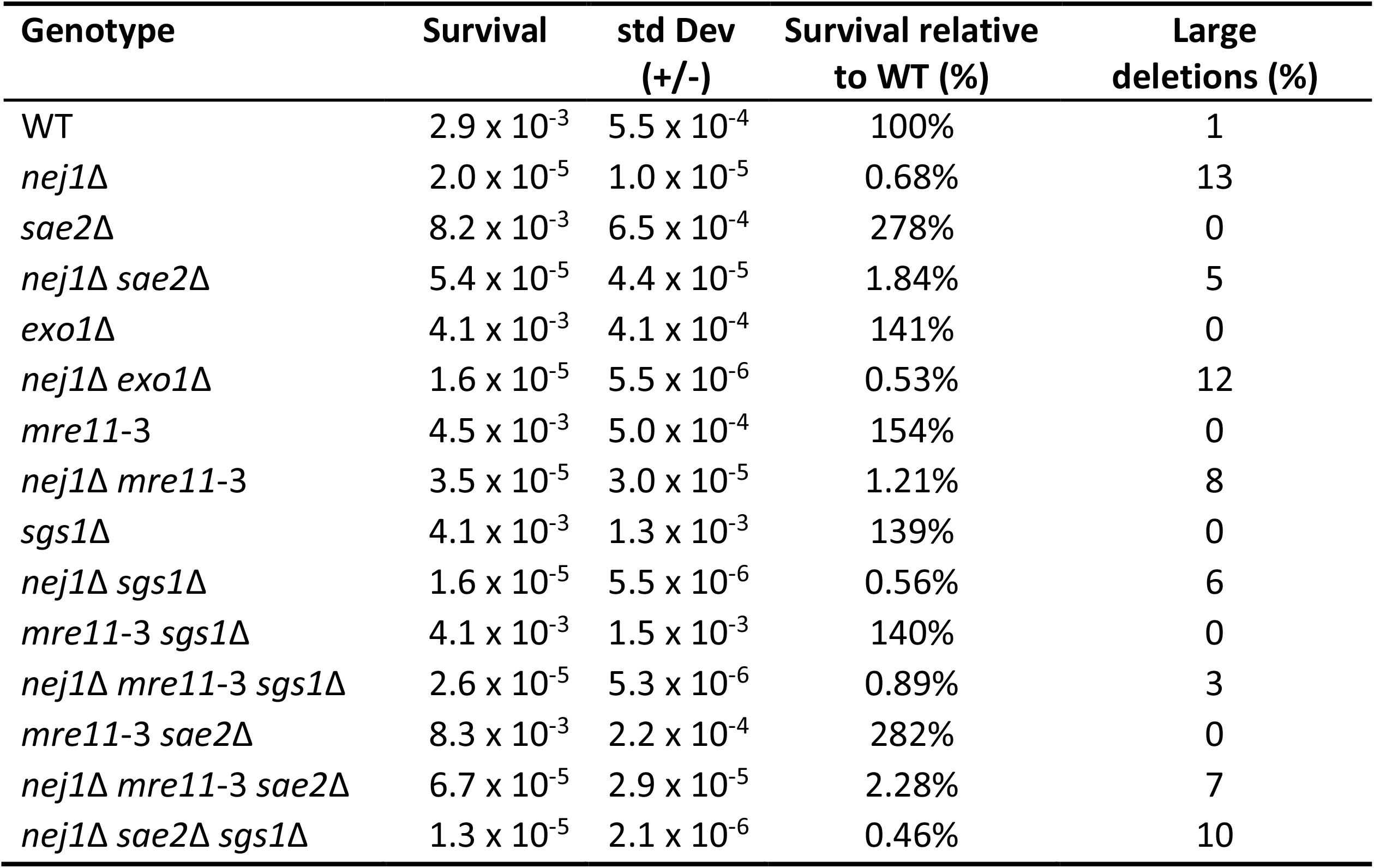
Survival and percentage of large deletions during continuous HO-induction.

We previously showed that large deletions at the DSB develop when DNA end-bridging is defective and 5’ resection proceeds [4]. Given that Sae2, like MRX and Nej1, also has a role in DNA end-bridging at DSBs [36-38], we wanted to assess this along with 5’ resection and determine how this correlated with the rate of genomic deletions that developed in the various *sae2*Δ mutant combinations. End-bridging was measured in cells where both sides of the DSB are tagged with fluorescent markers. The TetO array and the LacO array were integrated 3.2 Kb and 5.2Kb respectively from the DSB in cells expressing TetR^GFP^ and LacO^mCherry^ fusions, enabling us to visualize both sides by fluorescence microscopy (Fig S4B). In asynchronous cells, the distance between the GFP and mCherry foci was measured 2 hrs after DSB induction. In wild type cells, the mean distance between the fluorescent markers was not significantly different after HO cutting (0.28 µm) compared to before galactose induction (0.24 µm), indicating successful DNA end-bridging at the break (Fig 3E). The distance between markers increased in *sae2Δ* mutant cells (0.44 µm), indicating a defect in end-bridging (Fig 3E). However, no significant increase in the distance was observed after DSB induction in *sgs1*Δ or *exo1*Δ mutants (Fig S4B). Deletion of *SAE2* and *NEJ1* showed an epistatic relationship, as the defect in double mutant cells was comparable to that of the single mutants (Fig 3E). 5’ resection increased in *sae2*Δ *nej1*Δ compared to *sae2*Δ, and this correlated with the development of large deletions in the double mutants. Collectively, our data support a role for Sae2 in end-bridging together with Nej1 and a role for Sae2 in 5’ resection together with Dna2 recruitment at DSBs, a pathway which is inhibited by Nej1 and distinct of Mre11 activation by Sae2.

### 3.4 *NEJ1* alleviates the synthetic lethality of *sae2*Δ *sgs1*Δ

We observed that recruitment of Dna2 to DSBs was partially dependent on Sae2 and Sgs1, and both pathways were inhibited by Nej1, prompting us to determine whether deletion of *NEJ1* would alleviate *sae2*Δ *sgs1*Δ synthetic lethality (SL) [28]. We crossed *sae2*Δ *nej1*Δ with *sgs1*Δ *nej1*Δ and spores with the triple mutant combination grew remarkably well (Fig 4A). The triple mutants showed reduced survival under conditions of continuous HO-DSB induction similar to other mutant combination containing *nej1*Δ and the frequency of large deletions that developed in *sae2*Δ *sgs1*Δ *nej1*Δ survivors was slightly above that of s*ae2*Δ *nej1*Δ or *sgs1*Δ *nej1*Δ double mutant survivors (Table 1 and Fig. S4A). The sensitivity of *sae2*Δ *sgs1*Δ *nej1*Δ mutants to Phleomycin and MMS was however similar in *sae2*Δ *nej1*Δ and *sgs1*Δ *nej1*Δ double mutant cells (Fig 4B).

**Fig 4.**
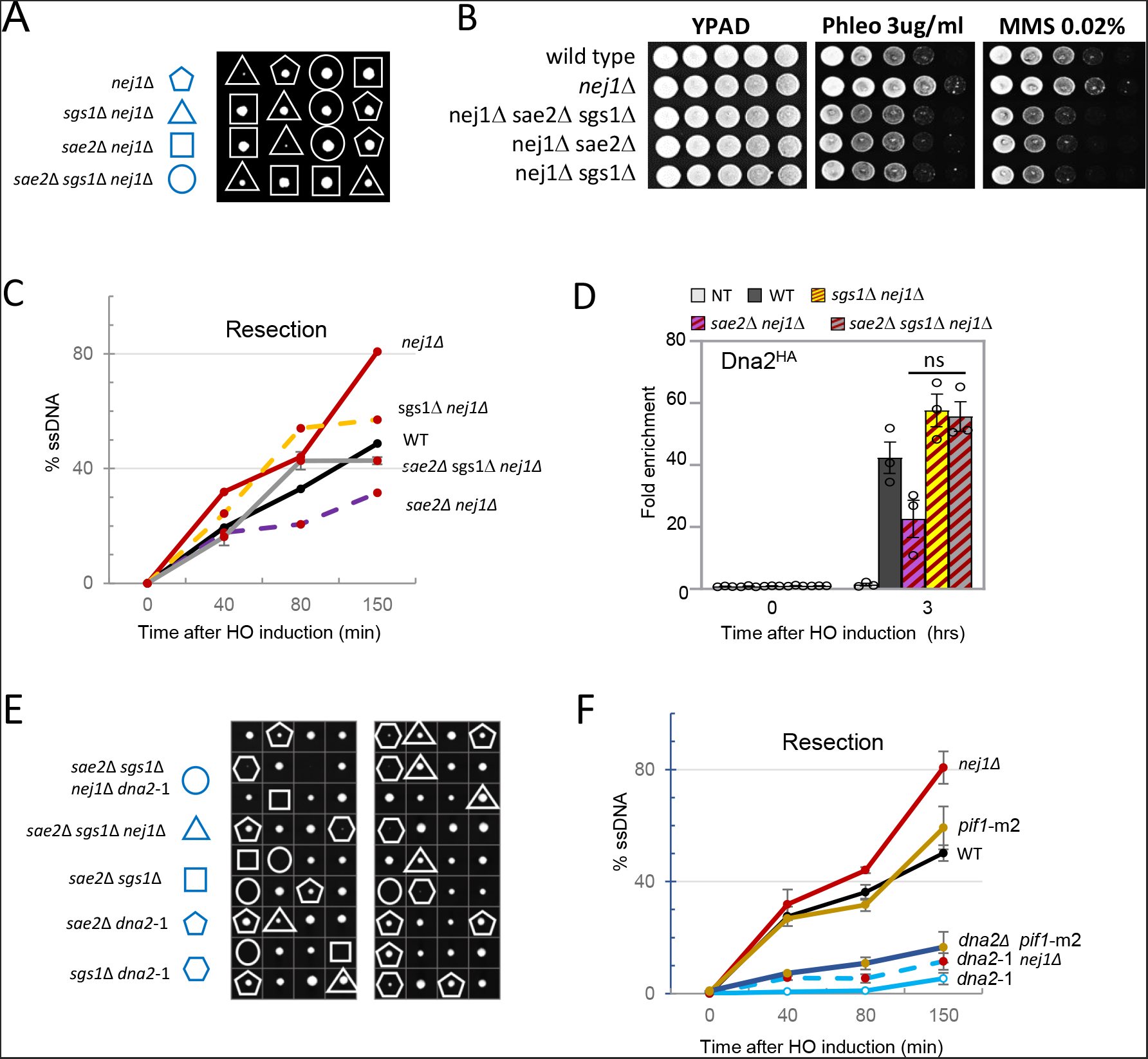
*NEJ1* alleviates the synthetic lethality of *sae2*Δ *sgs1*Δ. **(A)** Viability and genotypes of spores derived from diploids of *sae2*Δ *nej1*Δ (JC-5675) and *sgs1*Δ *nej1*Δ (JC-3885). **(B)** Five-fold serial dilutions of the cells in wild type (JC-727), *nej1*Δ (JC-1342), *sae2*Δ *sgs1*Δ *nej1*Δ (JC-5750), *sae2*Δ *nej1*Δ (JC-5675), *sgs1*Δ *nej1*Δ (JC-3759) were spotted on YPAD, Phleomycin 3.0 ug/ml, and MMS 0.02%. **(C)** 5’ DNA Resection 0.15kb away from the HO DSB using a qPCR-based approach described in the methods section. Frequency of resection is plotted as % ssDNA at 0, 40, 80 and 150 mins. post DSB induction in cycling cells in wild type (JC-727), *nej1*Δ (JC-1342), *sae2*Δ *nej1*Δ (JC-5675), *sgs1*Δ *nej1*Δ (JC-3759) and *sae2*Δ *sgs1*Δ *nej1*Δ (JC-5750). **(D)** Enrichment of Dna2^HA^ at 0.6kb from DSB 0 and 3 hours after DSB induction in wild type (JC-4117), *sae2*Δ *nej1*Δ (JC-5597), *sgs1*Δ *nej1*Δ (JC-5627), *sae2*Δ *sgs1*Δ *nej1*Δ (JC-5480) and no tag control (JC-727) was determined. The fold enrichment is normalized to recovery at the SMC2 locus. **(E)** Viability and genotypes of spores derived from heterozygous diploids of *SAE2*+/*sae2*Δ, *SGS1+/sgs1*Δ, *NEJ1*+/ *nej1*Δ and *DNA2*+/*dna2*-1 generated from a cross between JC-5749 and JC-5655. **(F)** 5’ DNA Resection 0.15kb away from the HO DSB using a qPCR-based approach described in the methods section. Frequency of resection is plotted as % ssDNA at 0, 40, 80 and 150 mins. post DSB induction in cycling cells in wild type (JC-727), *nej1*Δ (JC-1342), *dna2*-1 (JC-5655), *dna2*-1 *nej1*Δ (JC-5670), *pif1*-m2 (yWH0056) and *dna2*Δ *pif1*-m2 (yWH0055). The *pif1*-m2 and *dna2*Δ *pif1*-m2 strains in the same background were a kind gift from Greg Ira’s laboratory, Baylor College of Medicine. For all the experiments - error bars represent the standard error of three replicates.

Furthermore, resection in *sae2*Δ *sgs1*Δ *nej1*Δ triple mutants was similar to wild type and significantly higher than in *sae2*Δ and *sgs1*Δ single mutants (Fig 4C, 1G, 2D). Moreover, Dna2 recovery at the DSB in *sae2*Δ *sgs1*Δ *nej1*Δ mutants was similar to *sgs1*Δ *nej1*Δ and higher than in *sgs1*Δ single mutants or *sae2*Δ +/-*NEJ1* (Figs 4D, 2C, 3A).

To determine whether suppression of *sae2*Δ *sgs1*Δ lethality by *NEJ1* deletion required Dna2 nuclease activity, we generated heterozygous diploids for *sae2*Δ *sgs1*Δ *nej1*Δ and nuclease deficient *dna2-1* (P504→S) and upon tetrad dissection recovered no viable spores with quadruple mutant combination (Fig 4E). By contrast, *sae2*Δ *sgs1*Δ *nej1*Δ *exo1*Δ spores were viable, thus suppression of *sae2*Δ *sgs1*Δ synthetic lethality by *nej1*Δ depends on the nuclease activity of Dna2, not Exo1 (Fig. S5A). Resection in *dna2*-1 and *dna2*Δ *pif1*-m2 was similar to each other and more defective than resection in *sgs1*Δ mutants, but similar to the resection defect seen in *sae2*Δ mutants (Fig. 4F and Fig. S5B). Taken together, these data underscore the importance of Sae2-Dna2 interactions and Dna2 nuclease activity in DSB repair.

## 4. Discussion

We suggest that Nej1 operates as a general inhibitor of 5’ resection at DSBs. Not only does Nej1 inhibit Dna2 interactions with Sgs1 and MRX [4], but it physically interacts with Sae2, inhibiting both MRX-dependent recruitment of Sae2 and Sae2-dependent recruitment of Dna2 to the DSB. Our data support a model whereby Dna2 is recruited to a DSB through three pathways, all of which are inhibited by Nej1 (Fig 5; panel a). Dna2 localizes primarily through binding with Sgs1 or Sae2, thus deleting both results in lethality (Fig 5; panel b). Removal of Nej1, which binds Mre11-C, allows Dna2 recruitment through Mre11-Dna2 interactions, suppressing *sae2*Δ *sgs1*Δ synthetic lethality [4, 28]. Sae2 can initiate resection through Mre11 activation, but in the absence of Mre11 nuclease activity and Sgs1 helicase, it can initiate resection through interactions with Dna2. Our data show that Sae2 can compensate for *sgs1*Δ to localize Dna2 to DSBs, however if both *SAE2* and *SGS1* are deleted, then Dna2 can localize through binding Mre11 if *NEJ1* is also deleted [4]. Consistent with this model, viability of *sae2*Δ *sgs1*Δ *nej1*Δ triple mutant depends on the nuclease activity of Dna2. After resection initiates, and Ku dissociates, Exo1 would serve as the nuclease in long-range resection (Fig 5; panel c).

**Fig 5.**
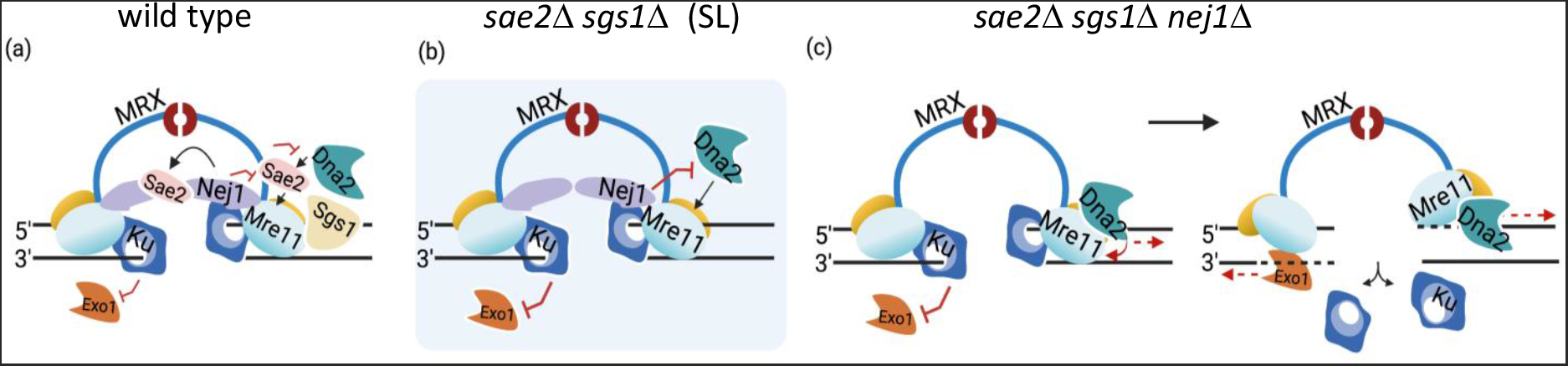
Interplay of Nej1, Sae2 and Sgs1 at DSB. Model of DSB where Nej1 prevents Mre11-dependent Dna2 recruitment to DSB. (a) In wild type cells Nej1 inhibits Dna2 recruitment via Mre11, Sae2 and Sgs1. (b) In *sae2*Δ *sgs1*Δ mutant cells Nej1 inhibits Dna2-Mre11 interaction and therefore prevents the residual Dna2 recruitment and resection, resulting into the synthetic lethality. (c) Upon *NEJ1* deletion, Dna2 can get recruited through Mre11 leading to resection and repair, resulting into alleviation of the synthetic lethality and growth of *sae2*Δ *sgs1*Δ *nej1*Δ cells. Created with BioRender.com

### 4.1. Sae2-dependent recruitment of Dna2 is inhibited by Nej1

Dna2 localization to DSBs is partly, but not entirely, dependent on Sgs1 helicase (Fig 2A,C). An alternative mode of Dna2 recruitment involves Sae2 (Fig 3A, [19]). Our results support previous work showing a role for human CtIP in Dna2 recruitment to DSBs [33], and provide mechanistic insight for *in vitro* studies where CtIP stimulates Dna2 nuclease [34, 35]. Although our findings differ slightly from previous work, which showed little decrease in Dna2 recovery 3 hours after DSB induction, the discrepancy could stem from slight variations in experimental design because in the same study Dna2 was reduced 2-hours after DSB induction in *sae2*Δ mutants [19].

Dna2 recovery was reduced more in *sae2*Δ than in *sgs1*Δ single mutants and upon further deletion of *NEJ1* were similar to wild type (Figs. 2C and 3A). In contrast to *sae2*Δ *nej1*Δ, Dna2 enrichment was above wild type in *sgs1*Δ *nej1*Δ mutants, and similar to *nej1*Δ, indicating that the Sgs1-independent pathway(s) of Dna2 localization were robustly inhibited by Nej1 (Fig 2C; [4]). Indeed, our data support this model as physical interactions between Sae2 and MRX, Sae2 and Dna2, and Mre11 and Dna2 increased in *nej1*Δ mutant cells (Fig 3C). Earlier work showed that the localization of MRX to DSBs was not disrupted in nuclease deficient *mre11*-3 mutants [12, 13], which is important as the MRX complex is needed for the recruitment of all the processing factors we investigated here. Furthermore, using *mre11*-3 we could also see that Dna2 recovery and resection trends were not significant affected by the disruption of Mre11 nuclease activity.

There was a marked decrease in resection in *sae2*Δ compared to *mre11*-3 mutants, which highlights and supports previous work showing that Sae2 has a role at DSBs in addition to Mre11 activation. Our data show that resection differences can be attributed to the decreased recovery of Dna2 in *sae2*Δ compared to *mre11*-3 mutants. Moreover, given the importance of Sae2 in Dna2 localization, resection in *mre11*-3 could even be supported by increased Sae2 levels in that mutant (Fig. 2E and [28]). These data do not contradict, but complement, earlier work showing that decreased resection resulted from increased end-protection by Ku in *sae2*Δ mutants [19]. Ku is important for Nej1 recruitment, therefore it is noteworthy that increased Ku did not result in increased Nej1 recovery in *sae2*Δ (Figs. 1D and 1SA). Furthermore, resection differences observed when comparing *sae2*Δ and *mre11*-3 mutants might also be related to checkpoint signalling defects in *sae2*Δ mutants, defects that are independent of Mre11 nuclease activity [28]. Lastly, our data do not contradict Sae2 functioning as a nuclease [29], however further studies involving Nej1, Dna2 with Sae2-mutants (D285P/K288P and E161P/K163P) would be needed to investigate this directly.

### 4.2. Nej1 and Sae2 in DNA end-bridging

Deletion of *NEJ1* and *SAE2* show epistatic end-bridging defects raising the possibility that Nej1 and Sae2 collaborate to restrain movement of the broken DNA ends at DSBs in contrast to their antagonistic roles in resection. Given the physical interaction between Nej1 and Sae2 (Fig 1E), it is possible that the two factors could function as a complex in end bridging. It might be informative to determine whether Sae2 and Nej1 can form hetero-oligomers, distinguishing a sub-population of Sae2 involved in DNA end-bridging apart from Sae2 homo-oligomers involved in Mre11 activation and checkpoint signalling [46, 47]. Importantly, disruptions in other factors involved in 5’ resection including *exo1*Δ, or *sgs1*Δ, did not show end-bridging defects (Fig S3). Furthermore, DNA end-bridging was also maintained in *mre11*-3 mutants, which is in line with previous work showing that the structural integrity of the MRX complex, but not its nuclease activity, is important for bridging [4, 12, 13, 48]. Comparing end-bridging defects in *mre11*-3 and *sae2*Δ mutants +/-*NEJ1* indeed supports the model that large deletions develop when 5’ resection proceeds and end-bridging is disrupted. In *mre11*-3 mutants, 5’ resection proceeds but end-bridging was not disrupted, whereas in *sae2*Δ mutants, bridging was disrupted but 5’ resection was very low and neither single mutant showed large deletions (Table 1 and [12, 13, 36-38]). By contrast, large deletions formed when either mutant was combined with *nej1*Δ, although the frequency was lower compared to *nej1*Δ single mutants (Table 1).

### 4.3 Synthetic lethality of *sae2*Δ *sgs1*Δ is supressed by *NEJ1* deletion

Suppression of *sae2*Δ *sgs1*Δ synthetic lethality by *nej1*Δ was dependent on Dna2, but not Exo1 nuclease activity (Fig 4F and S4) [28]. Moreover, the higher rate of resection in *sgs1*Δ mutants compared to *dna2*Δ *pif1*-m2 and *dna2*-1 mutants also demonstrates the importance of Dna2 in DSB repair, independently of Sgs1. While both Dna2 and Sgs1 have important links to the DNA damage checkpoint [49], the greater resection defect in *dna2*-1 is likely not attributed to its checkpoint functions as mutations in Dna2 that disrupt signalling map to its N-terminal region, distinct of its nuclease and helicase activities [50]. In addition, 5’ resection was similarly reduced in *dna2*-1 and *dna2*Δ *pif1*-m2 mutants (Fig 4D), ruling out a potential dominant-negative effect for *dna2*-1 in tetrad analysis.

Surprisingly, the frequency of 5’ resection and the recovery level of Dna2 in *sae2*Δ *sgs1*Δ *nej1*Δ triple mutants was similar to wild type, and above *sae2*Δ *nej1*Δ (Fig 4C, D), suggesting that Sgs1 is inhibitory in *sae2*Δ *nej1*Δ double mutant cells. We previously showed that both Sgs1 and Dna2 interact directly with Mre11 [4], thus in *sae2*Δ *nej1*Δ mutants the presence of Sgs1 could inhibit the initiation of resection occurring from Dna2-Mre11 interactions. The presence of Sgs1, and therefore Dna2-Sgs1 complex formation, might be less efficient at initiating resection compared to its abilities in long-range resection. Like with *nej1*Δ, previous work showed that *ku70*Δ and *rad9*Δ also suppressed *sae2*Δ *sgs1*Δ lethality [20, 28]. This raises the possibility that suppression of *sae2*Δ *sgs1*Δ lethality might result from a decrease in overall NHEJ when *NEJ1* was also deleted. However, two results argue that intrinsic loss of NHEJ itself does not suppress this lethality. First, deletion of *DNL4* ligase does not rescue *sae2*Δ *sgs1*Δ and second, NHEJ occurs in *sae2*Δ *sgs1*Δ *rad9*Δ triple mutants [20, 28]. Taken together, our work provides new information on how Nej1 inhibits nuclease recruitment and 5’ resection at DSBs. These functions help preserves genome integrity during repair pathway choice and ascribe a wider-range of responsibilities to Nej1 that are distinct of its roles in canonical NHEJ.

## 2. Materials and Methods

All the yeast strains used in this study are listed in Table S3 and were obtained by crosses. The strains were grown on various media in experiments described below. For HO-induction of a DSB, YPLG media is used (1% yeast extract, 2% bactopeptone, 2% lactic acid, 3% glycerol and 0.05% glucose). For the continuous DSB assay, YPA plates are used (1% yeast extract, 2% bacto peptone, 0.0025% adenine) supplemented with either 2% glucose or 2% galactose. For the mating type assays, YPAD plates are used (1% yeast extract, 2% bacto peptone, 0.0025% adenine, 2% dextrose). For yeast 2-hybrid assays, standard amino acid drop-out media lacking histidine, tryptophan and uracil is used and 2% raffinose is added as the carbon source for the cells.

### 2.1. Chromatin Immunoprecipitation

ChIP assay was performed as described previously [4]. Cells were cultured overnight in YPLG at 25°C. Cells were then diluted to 5 × 10^6^ cells/ml and were cultured to one doubling (3-4 hrs) at 30°C. 2% GAL was added to the YPLG and cells were harvested and crosslinked at various time points using 3.7% formaldehyde solution. Following crosslinking, the cells were washed with ice cold PBS and the pellet stored at -80°C. The pellet was re-suspended in lysis buffer (50mM Hepes pH 7.5, 1mM EDTA, 80mM NaCl, 1% Triton, 1mM PMSF and protease inhibitor cocktail) and cells were lysed using Zirconia beads and a bead beater. Chromatin fractionation was performed to enhance the chromatin bound nuclear fraction by spinning the cell lysate at 13,200rpm for 15 minutes and discarding the supernatant. The pellet was re-suspended in lysis buffer and sonicated to yield DNA fragments (∼500bps in length). The sonicated lysate was then incubated in beads + anti-HA/Myc Antibody or unconjugated beads (control) for 2 hrs at 4°C. The beads were washed using wash buffer (100mM Tris pH 8, 250mM LiCl, 150mM (HA Ab) or 500mM (Myc Ab) NaCl, 0.5% NP-40, 1mM EDTA, 1mM PMSF and protease inhibitor cocktail) and protein-DNA complex was eluted by reverse crosslinking using 1%SDS in TE buffer, followed by proteinase K treatment and DNA isolation via phenol-chloroform-isoamyl alcohol extraction. Quantitative PCR was performed using the Applied Biosystem QuantStudio 6 Flex machine. PerfeCTa qPCR SuperMix, ROX was used to visualize enrichment at HO2 (0.5kb from DSB) and HO1 (1.6kb from DSB) and SMC2 was used as an internal control. HO-cutting was measured in strains used to perform ChIP in Table S2.

### 2.2. Tethering Microscopy

Cells derived from the parent strain JC-4066 were diluted and grown overnight in YPLG at 25°C to reach a concentration of 1×10^7^ cells/ml. Cells were treated with 2% GAL for 2 hours and cell pellets were collected and washed 2 times with PBS. After the final wash, cells were placed on cover slips and imaged using a fully motorized Nikon Ti Eclipse inverted epi-fluorescence microscope. Z-stack images were acquired with 200 nm increments along the z plane, using a 60X oil immersion 1.4 N.A. objective. Images were captured with a Hamamatsu Orca flash 4.0 v2 sCMOS 16-bit camera and the system was controlled by Nikon NIS-Element Imaging Software (Version 5.00). All images were deconvolved with Huygens Essential version 18.10 (Scientific Volume Imaging, The Netherlands, http://svi.nl), using the Classic Maximum Likelihood Estimation (CMLE) algorithm, with SNR:40 and 50 iterations. To measure the distance between the GFP and mCherry foci, the ImageJ plug-in Distance Analysis (DiAna) was used [51]. Distance measurements represent the shortest distance between the brightest pixel in the mCherry channel and the GFP channel. Each cell was measured individually and > 50 cells were analyzed per condition per biological replicate.

### 2.3. qPCR based Resection Assay

Cells from each strain were grown overnight in 15ml YPLG to reach an exponentially growing culture of 1×10^7^ cells/mL. Next, 2.5mL of the cells were pelleted as timepoint 0 sample, and 2% GAL was added to the remaining cells, to induce a DSB. Following that, respective timepoint samples were collected. Genomic DNA was purified using standard genomic preparation method by isopropanol precipitation and ethanol washing, and DNA was re-suspended in 100mL ddH_2_O. Genomic DNA was treated with 0.005μg/μL RNase A for 45min at 37°C. 2μL of DNA was added to tubes containing CutSmart buffer with or without RsaI restriction enzyme and incubated at 37°C for 2hrs. Quantitative PCR was performed using the Applied Biosystem QuantStudio 6 Flex machine. PowerUp SYBR Green Master Mix was used to quantify resection at MAT1 (0.15kb from DSB) locus. Pre1 was used as a negative control. % resected / cut HO loci is reported from the amount of RsaI cut DNA normalized to the level of HO cutting at each timepoint (Table S1) [36].

### 2.4. Continuous DSB assay and identification of mutations in survivors

Cells were grown overnight in YPLG media at 25°C to saturation. Cells were collected by centrifugation at 2500rpm for 3 minutes and pellets were washed 1x in ddH_2_O and re-suspended in ddH_2_O. Cells were counted and spread on YPA plates supplemented with either 2% GLU or 2% GAL. On the Glucose plates 1×10^3^ total cells were added and on the galactose plates 1×10^5^ total cells were added. The cells were incubated for 3-4 days at room temperature and colonies counted on each plate. Survival was determined by normalizing the number of surviving colonies in the GAL plates to number of colonies in the GLU plates. 100 survivors from each strain were scored for the mating type assay as previously described [9].

### 2.5. Yeast 2-hybrid

Various plasmids (Table S4) were constructed containing the gene encoding the region of the proteins – Sae2, Dna2, Mre11, Nej1, Rad50 and Xrs2, using the primers listed in Table S5. The plasmids J-965 and J-1493 and the inserts were treated with BamHI and EcoRI and ligated using T4 DNA ligase. The plasmids were sequence verified. Reporter (J-359), bait (J-965) and prey (J-1493) plasmids, containing the gene encoding the desired protein under a galactose inducible promoter, were transformed into JC-1280. Cells were grown overnight in –URA –HIS –TRP media with 2% raffinose. Next day, cells were transferred into –URA –HIS –TRP media with either 2% GLU or 2% GAL and grown for 6 hrs at 30°C. Cell pellets were resuspended and then permeabilized using 0.1% SDS followed by ONPG addition. β-galactosidase activity was estimated by measuring the OD at 420nm, relative β-galactosidase units were determined by normalizing to total cell density at OD600. Additionally, for drop assay cells were grown and spotted in five-fold serial dilutions on plates containing 2% galactose lacking histidine and tryptophan (for plasmid selection) and leucine (for measuring expression from *lexAop6-LEU2*). Plates were photographed after 3-4 days of incubation at 30°C.

### 2.6. Western Blots

Cells were lysed by re-suspending them in lysis buffer (with PMSF and protease inhibitor cocktail tablets) followed by bead beating. The protein concentration of the whole cell extract was determined using the NanoDrop. Equal amounts of whole cell extract were added to wells of 10% polyacrylamide SDS gel. After the run, the protein were transferred to Nitrocellulose membrane at 100V for 80mins. The membrane was Ponceau stained (which served as a loading control), followed by blocking in 10% milk-PBST for 1hour at room temperature. Respective primary antibody solution (1:1000 dilution) was added and left for overnight incubation at 4°C. The membranes were then washed with PBST and left for 1 hour with secondary antibody. Followed by washing the membranes, adding the ECL substrates and imaging them.

### 2.7. Tetrad Analysis

Diploids of *sae2*Δ *nej1*Δ (JC-5675) X *sgs1*Δ *nej1*Δ (JC-3885) (Figure 4A); and *sae2*Δ *sgs1*Δ *nej1*Δ (JC-5749) X *dna2*-1 (JC-5655) (Figure 4E) were sporulated. The spores (*sae2*Δ::HIS, *sgs1*Δ::NAT *nej1*Δ::KAN, *dna2*-1::TS) were checked by replica-plating on the marker plates (-HIS, +NAT, +KAN and 37°C). For analysis two-two gene segregation was observed among the tetrads. The tetrad scoring data is available with the paper.

### 2.8. DSB Efficiency

The efficiency of HO-cutting was measured as previously described at all timepoints in the 5’ resection experiments [9]. Cells were grown in YPLG before the addition of galactose to induce expression of the HO-endonuclease, leading to DSB formation. The cells were pelleted and gDNA was prepared followed by qPCR with a primer set flanking the DSB (HO6 primers, Table S5).

## Supporting information

**S1 Fig.**
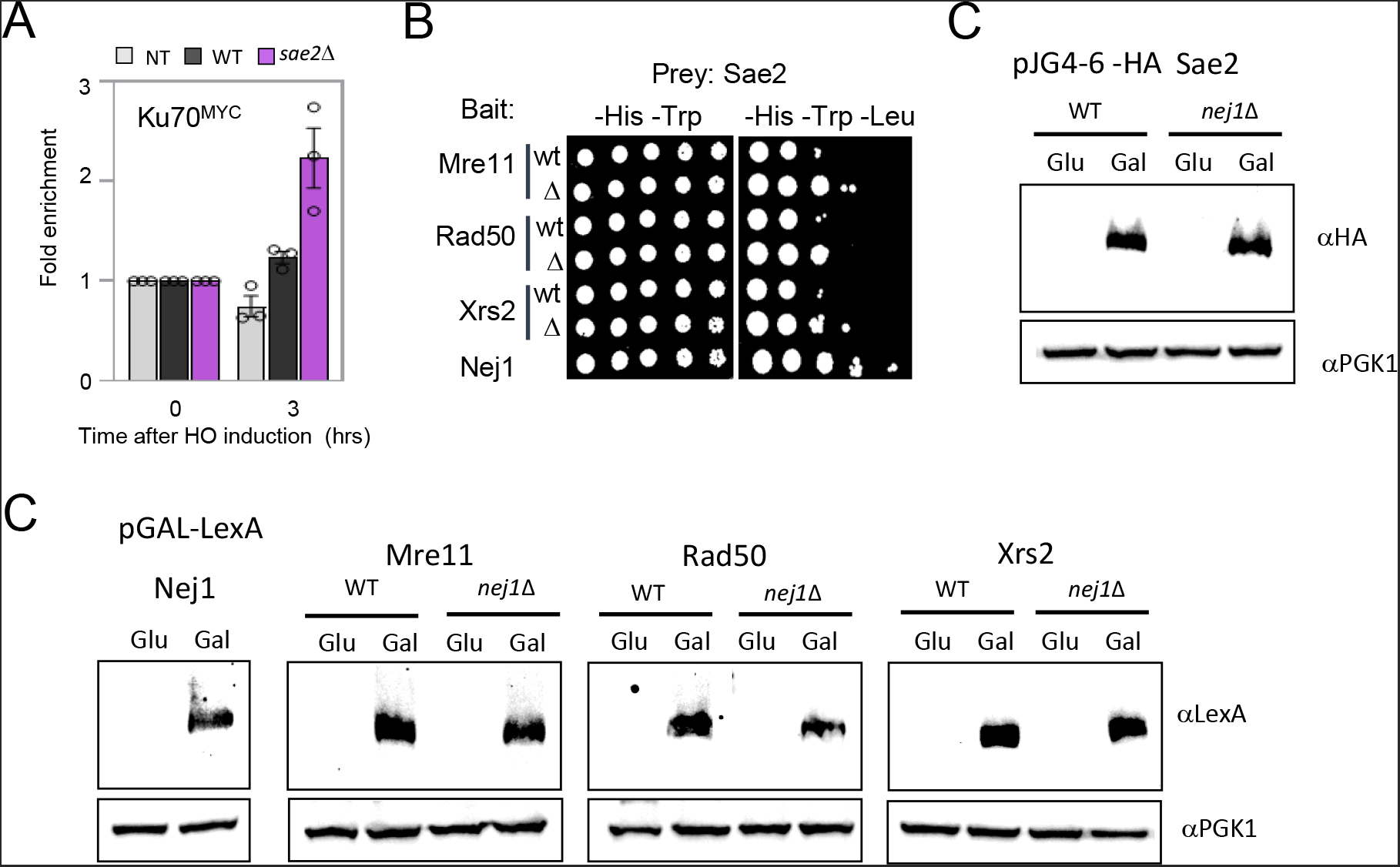
Ku recovery when *SAE2* is deleted and supporting data for Yeast 2 Hybrid analysis. **(A)** Enrichment of Ku70^Myc^ at DSB, at 0 and 3 hours, in wild type (JC-2271), *sae2*Δ (JC-5534) and no tag control (JC-727). The fold enrichment represents normalization over the SMC2 locus. **(B)** Y2H analysis of Sae2 fused to HA-AD, and Mre11, Rad50, Xrs2, or Nej1 fused to LexA-DBD was performed in wild type cells (JC-1280) and in isogenic cells with *nej1*Δ (JC-4556) using a drop assay on drop-out (-His, -Trp, -Leu) selective media plates. **(C-D)** Expression of proteins upon galactose induction. Western blots showing similar expression for HA-and LexA-tagged fusion proteins upon galactose induction in WT and *nej1*Δ. PGK1 was stained as a loading control.

**S2 Fig.**
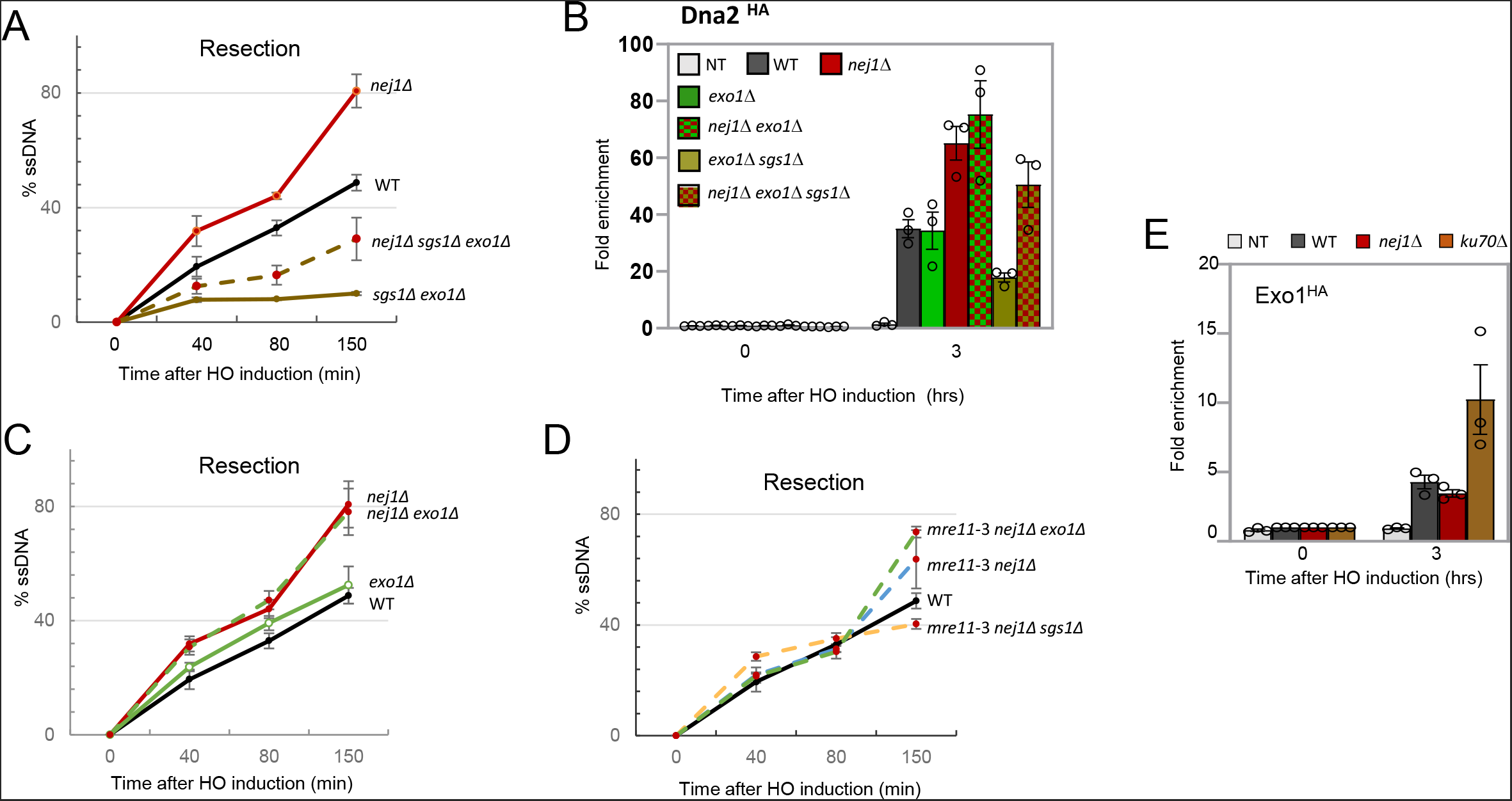
Dna2 recruitment is Exo1-independent. **(A, C and D)** 5’ DNA Resection 0.15kb away from the HO DSB using a qPCR-based approach described in the methods section. Frequency of resection is plotted as % ssDNA at 0, 40, 80 and 150 mins. post DSB induction in cycling cells in wild type (JC-727), *nej1*Δ (JC-1342), *exo1*Δ *sgs1*Δ (JC-4520), *exo1*Δ *sgs1*Δ *nej1*Δ (JC-5493), *exo1*Δ (JC-3767), *exo1*Δ *nej1*Δ (JC-5373), *mre11*-3 *nej1*Δ (JC-5369), *mre11*-3 *nej1*Δ *exo1*Δ (JC-5371) and *mre11*-3 *nej1*Δ *sgs1*Δ (JC-5667). **(B)** Enrichment of Dna2^HA^ at DSB, at 0 and 3 hours, in wild type (JC-4117), *nej1*Δ (JC-4118), *exo1*Δ (JC-5626), *exo1*Δ *nej1*Δ (JC-5666), *exo1*Δ *sgs1*Δ (JC-5559), *exo1*Δ *sgs1*Δ *nej1*Δ (JC-5566) and no tag control (JC-727). **(E)** Enrichment of Exo1^HA^ at DSB, at 0 and 3 hours, in wild type (JC-4869), *nej1*Δ (JC-6023), *ku70*Δ (JC-6018) and no tag control (JC-727). All the ChIP signals were determined at 0.6kb from DSB. The fold enrichment is normalized to recovery at the SMC2 locus. For all the experiments - error bars represent the standard error of three replicates.

**S3 Fig.**
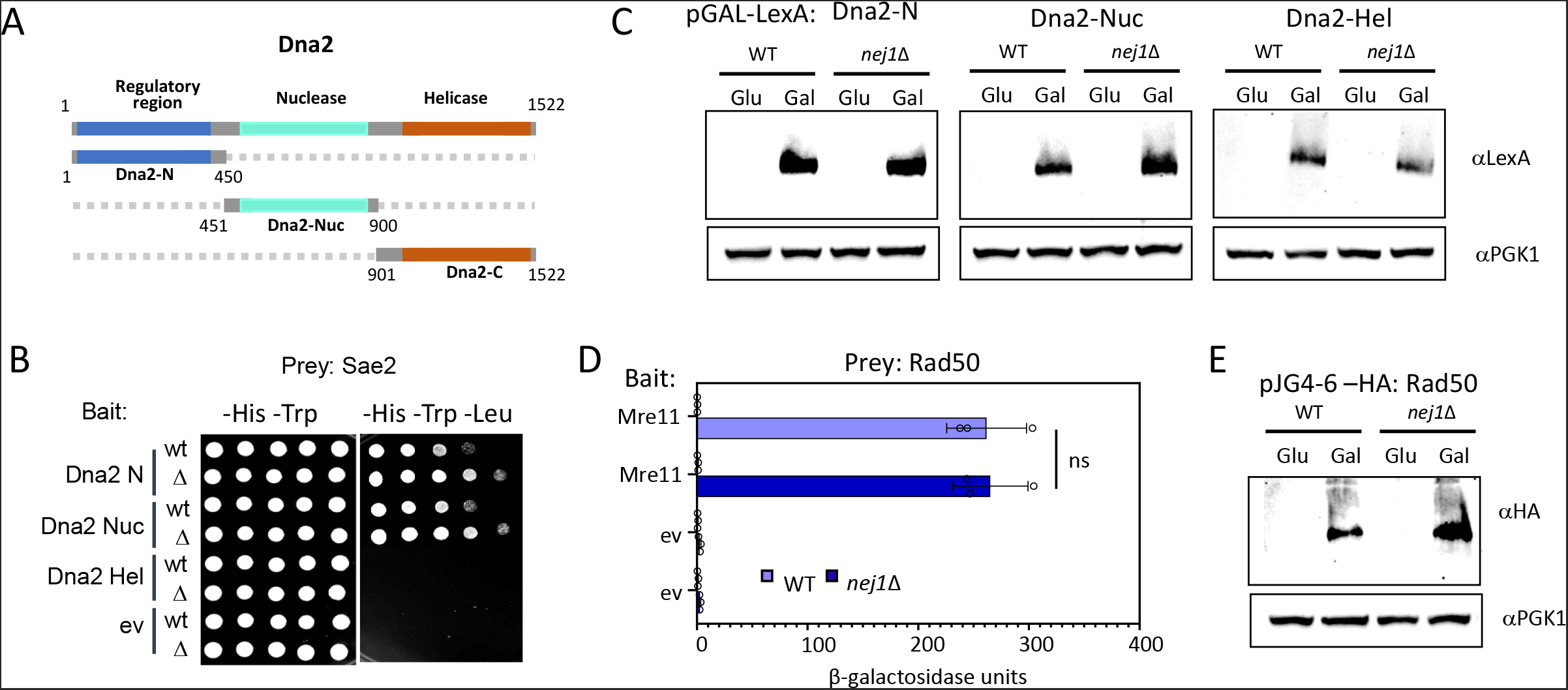
Supporting data for Yeast 2 Hybrid analysis. **(A)** Schematic representation of Dna2 and its functional domains. In dark blue is the N-terminal region, in light blue is the nuclease domain, and in orange is in the C-terminal helicase domain. **(B)** Y2H analysis of Sae2 fused to HA-AD, and Dna2 domains Nej1 fused to LexA-DBD was performed in wild type cells (JC-1280) and in isogenic cells with *nej1*Δ (JC-4556) using a drop assay on drop-out (-His, -Trp, -Leu) selective media plates. **(C and E)** Expression of proteins upon galactose induction. Western blots showing similar expression for LexA- and HA-tagged fusion proteins upon galactose induction in WT and *nej1*Δ. PGK1 was stained as a loading control. **(D)** Y2H analysis of Rad50 fused to HA-AD and Mre11 fused to LexA-DBD was performed in wild type cells (JC-1280) and in isogenic cells with *nej1*Δ (JC-4556) using a quantitative β-galactosidase assay.

**S4 Fig.**
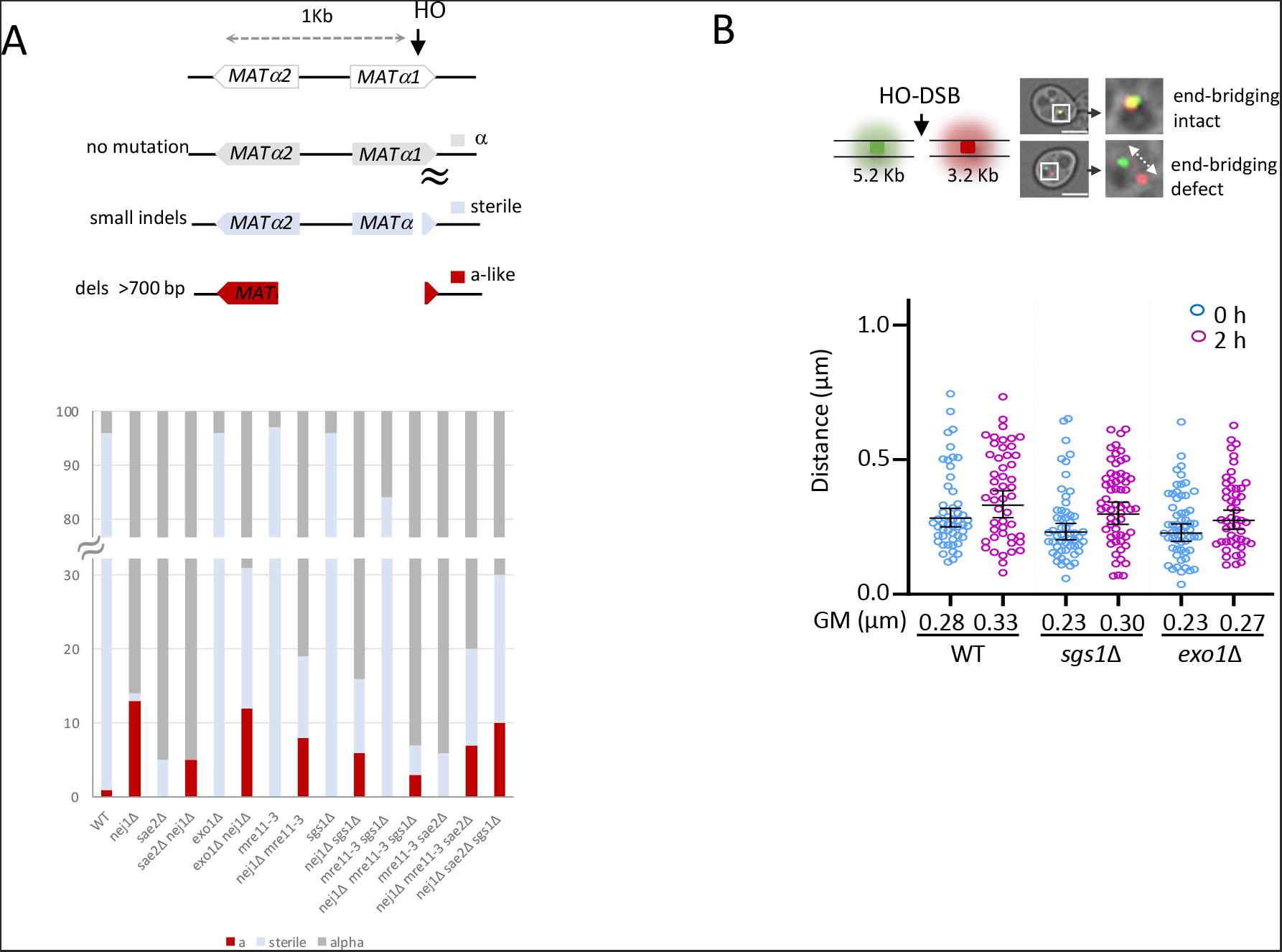
Microscopy and end-bridging measurements in various mutants. **(A) Above:** Schematic representation of mating type analysis of survivors from persistent DSB induction assays. The mating phenotype is a read out for the type of repair: α survivors (mutated HO endonuclease-site, grey), sterile survivors (small insertions and deletions, blue) and “a-like” survivors (>700 bps deletion, red). **Below:** Mating type analysis of survivors from persistent DSB induction assays in wild type (JC-727), *nej1*Δ (JC-1342), *sae2*Δ (JC-5673), *sae2*Δ *nej1*Δ (JC-5675), *exo1*Δ (JC-3767), *exo1*Δ *nej1*Δ (JC-5373),*mre11*-3 (JC-5372), *mre11*-3 *nej1*Δ (JC-5369), *sgs1*Δ (JC-3757), *sgs1*Δ *nej1*Δ (JC-3759), *mre11*-3 *sgs1*Δ (JC-5405), *mre11*-3 *sgs1*Δ *nej1*Δ (JC-5667) *mre11*-3 *sae2*Δ (JC-5501), *mre11*-3 *sae2*Δ *nej1*Δ (JC-5500) and *sae2*Δ *sgs1*Δ *nej1*Δ (JC-5750). **(B) Above:** Representative image of yeast cells with tethered (co-localized GFP and mCherry) and untethered (distant GFP and mCherry) ends. **Below:** Scatter data plot showing the tethering of DSB ends, at 0 and 2 hours, as measured by the distance between the GFP and mCherry foci in wild type (JC-4066), *sgs1*Δ (JC-5699) and *exo1*Δ (JC-5700). The Geometric mean (GM) distance for each sample is specified under respective sample data plot. Significance was determined using Kruskal-Wallis and Dunn’s multiple comparison test.

**S5 Fig.**
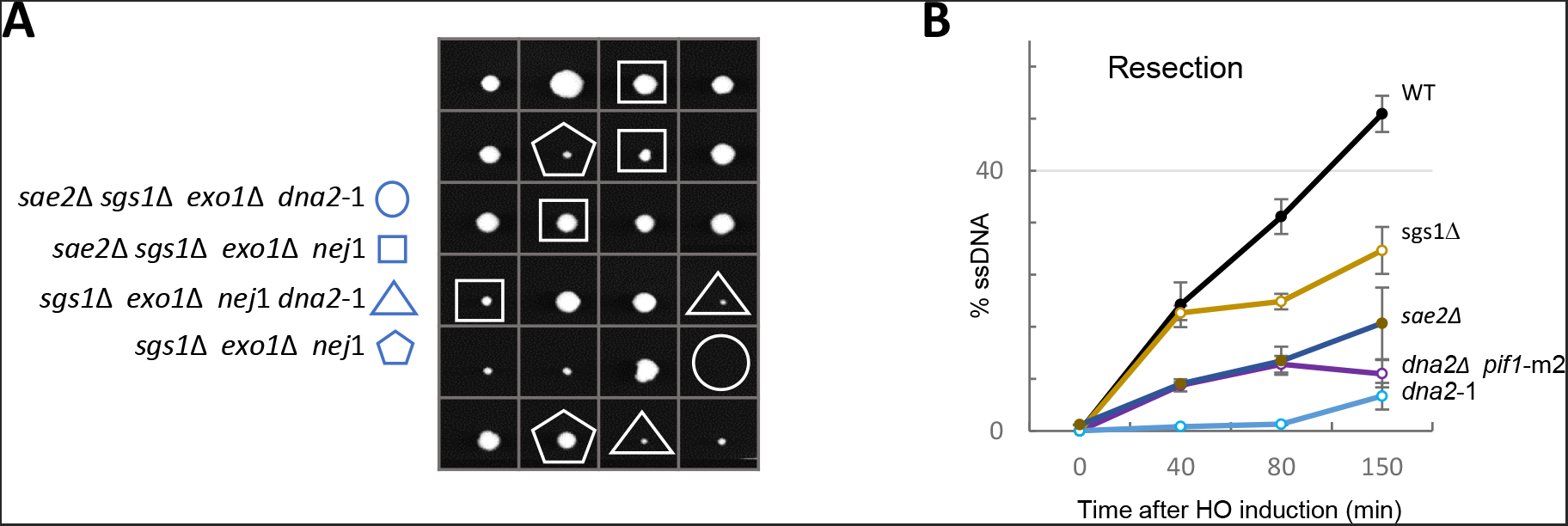
Tetrad analysis with Exo1. **(A)** Viability and genotypes of spores derived from diploids of *sae2*Δ *sgs1*Δ *nej1*Δ (JC-5749) and *exo1*Δ *dna2*-1 (JC-5692). The spores were checked by replica-plating on the marker plates (-HIS, +NAT, +KAN and 37°C). For analysis two-two gene segregation was observed among the tetrads. **(B)** 5’ DNA Resection 0.15kb away from the HO DSB using a qPCR-based approach described in the methods section. Frequency of resection is plotted as %ssDNA at 0, 40, 80 and 150 mins. post DSB induction in cycling cells in wild type (JC-727), *sae2*Δ (JC-5673), *sgs1*Δ (JC-3757), *dna2*-1 (JC-5655) and *dna2*Δ *pif1*-m2 (yWH0055). The *dna2*Δ *pif1*-m2 strains in the same background were a kind gift from Greg Ira’s laboratory, Baylor College of Medicine.

## Acknowledgments

We thank Dr. Greg Ira at Baylor College of Medicine for providing us with yeast mutants critical for the study. Our work was supported by operating grants from CIHR MOP-82736; MOP-137062 and NSERC 418122 awarded to J.A.C. The authors declare that they have no conflicts of interest with the contents of this article. We acknowledge the resources provided by the Live Cell Imaging Laboratory. The Nikon Ti Eclipse inverted epi-fluorescence microscope system was purchased with funds from the International Microbiome Centre, which is supported by the Cumming School of Medicine at University of Calgary, Western Economic Diversification (WED) and Alberta Economic Development and Trade (AEDT), Canada.

## Authors Contributions

A.M. and J.A.C. designed the research. A.M., and N.A. performed experiments and analyzed the data. A.M., N.A. and J.A.C wrote the manuscript.

## Supplemental Files

**S1 Table:** DSB cut efficiency in various mutants during resection measurements.

| Strain | Genotype | 40 min | 80 min | 150 min |
| --- | --- | --- | --- | --- |
| JC-727 | Wild type | 50.9 | 86.0 | 95.5 |
| JC-1342 | *nej1*Δ | 53.3 | 85.4 | 94.2 |
| JC-5673 | *sae2*Δ | 51.1 | 69.3 | 81.3 |
| JC-5675 | *sae2*Δ *nej1*Δ | 58.5 | 79.5 | 92.4 |
| JC-5372 | *mre11-3* | 75.2 | 88.8 | 93.1 |
| JC-5369 | *mre11-3 nej1*Δ | 80.1 | 90.8 | 91.9 |
| JC-3757 | *sgs1*Δ | 52.1 | 85.7 | 94.9 |
| JC-3759 | *sgs1*Δ *nej1*Δ | 55.3 | 77.3 | 89.5 |
| JC-5405 | *mre11-3 sgs1*Δ | 77.8 | 89.9 | 92.5 |
| JC-5667 | *mre11-3 sgs1*Δ *nej1*Δ | 63.2 | 68.9 | 84.3 |
| JC-5501 | *sae2*Δ *mre11-3* | 65.7 | 80.9 | 91.9 |
| JC-5500 | *sae2*Δ *mre11-3 nej1*Δ | 62.3 | 80.2 | 92.1 |
| JC-3767 | *exo1*Δ | 51.7 | 77.2 | 91.3 |
| JC-5373 | *exo1*Δ *nej1*Δ | 57.3 | 57.7 | 81.8 |
| JC-4520 | *exo1*Δ *sgs1*Δ | 66.0 | 73.7 | 74.9 |
| JC-5493 | *exo1*Δ *sgs1*Δ *nej1*Δ | 61.9 | 66.6 | 78.7 |
| JC-5750 | *sae2*Δ *sgs1*Δ *nej1*Δ | 44.4 | 76.0 | 84.5 |
| JC-5655 | *dna2-1* | 39.7 | 79.1 | 90.6 |
| JC-5670 | *dna2-1 nej1*Δ | 49.8 | 89.5 | 95.0 |
| yWH0056 | *pif1-m2* | 89.1 | 95.4 | 96.7 |
| yWH0055 | *dna2*Δ *pif1-m2* | 88.2 | 90.5 | 95.5 |

**S2 Table:** DSB cut efficiency in various mutants for ChIP.

| Strain | Genotype | HO Cutting (%)<br>at 180 min |
| --- | --- | --- |
| JC-4117 | Dna2<sup>HA</sup> | 80.33 |
| JC-4118 | Dna2<sup>HA</sup> *nej1*Δ | 87.05 |
| JC-5624 | Dna2<sup>HA</sup> *sgs1*Δ | 97.62 |
| JC-5594 | Dna2<sup>HA</sup> *mre11-3* | 98.14 |
| JC-5596 | Dna2<sup>HA</sup> *mre11-3 nej1*Δ | 89.02 |
| JC-5621 | Dna2<sup>HA</sup> *mre11-3 sgs1*Δ | 90.80 |
| JC-5623 | Dna2<sup>HA</sup> *mre11-3 sgs1*Δ *nej1*Δ | 76.04 |
| JC-5562 | Dna2<sup>HA</sup> *sae2*Δ | 81.14 |
| JC-5597 | Dna2<sup>HA</sup> *sae2*Δ *nej1*Δ | 87.73 |
| JC-5598 | Dna2<sup>HA</sup> *sae2*Δ *mre11-3* | 98.53 |
| JC-5593 | Dna2<sup>HA</sup> *sae2*Δ *mre11-3 nej1*Δ | 96.33 |
| JC-5480 | Dna2<sup>HA</sup> *sae2*Δ *sgs1*Δ *nej1*Δ | 91.34 |
| JC-5116 | Sae2<sup>HA</sup> | 85.93 |
| JC-5122 | Sae2<sup>HA</sup> *mre11*Δ | 88.73 |
| JC-5119 | Sae2<sup>HA</sup> *mre11-3* | 92.93 |
| JC-5124 | Sae2<sup>HA</sup> *nej1*Δ | 91.53 |
| JC-5702 | Sae2<sup>HA</sup> *mre11-3 nej1*Δ | 89.97 |

**S3 Table:**
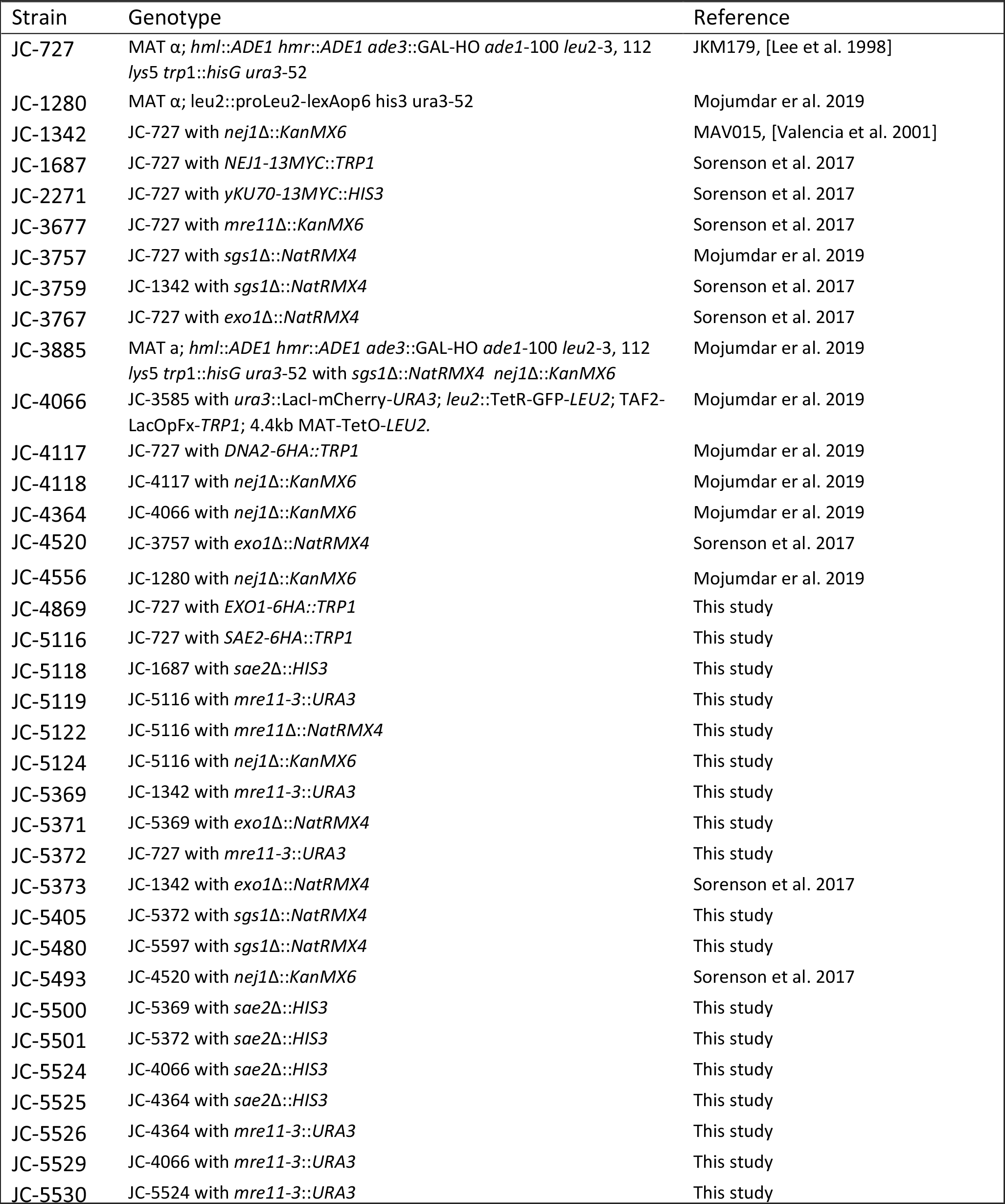
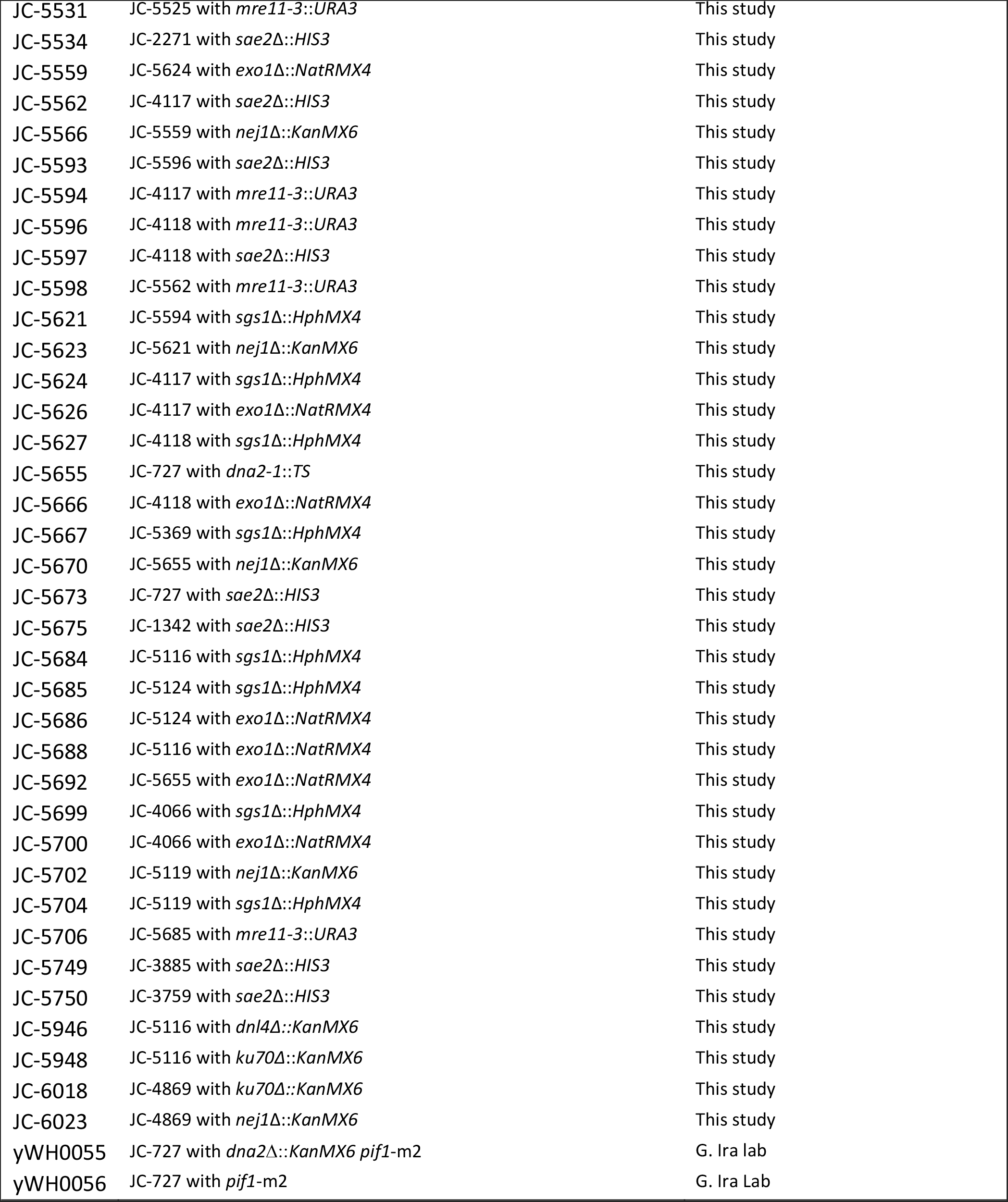
S. cerevisiae strains used in this study.

**S4 Table:** Plasmids used in this study.

| Plasmid number | Description |
| --- | --- |
| J-965 | pGAL-lexA |
| J-1493 | pJG4-6 |
| J-359 | pSH18-34 lexAGal1-lacZ |
| J-125 | J-965 with Nej1 |
| J-186 | J-965 with Mre11 |
| J-199 | J-965 with Xrs2 |
| J-200 | J-965 with Rad50 |
| J-1855 | J-965 with Dna2-(1-440) N-term |
| J-1856 | J-965 with Dna2-(441-920) Nuclease |
| J-1857 | J-965 with Dna2-(921-1522) Helicase |
| J-1898 | J-1493 with Sae2 |

**S5 Table:**
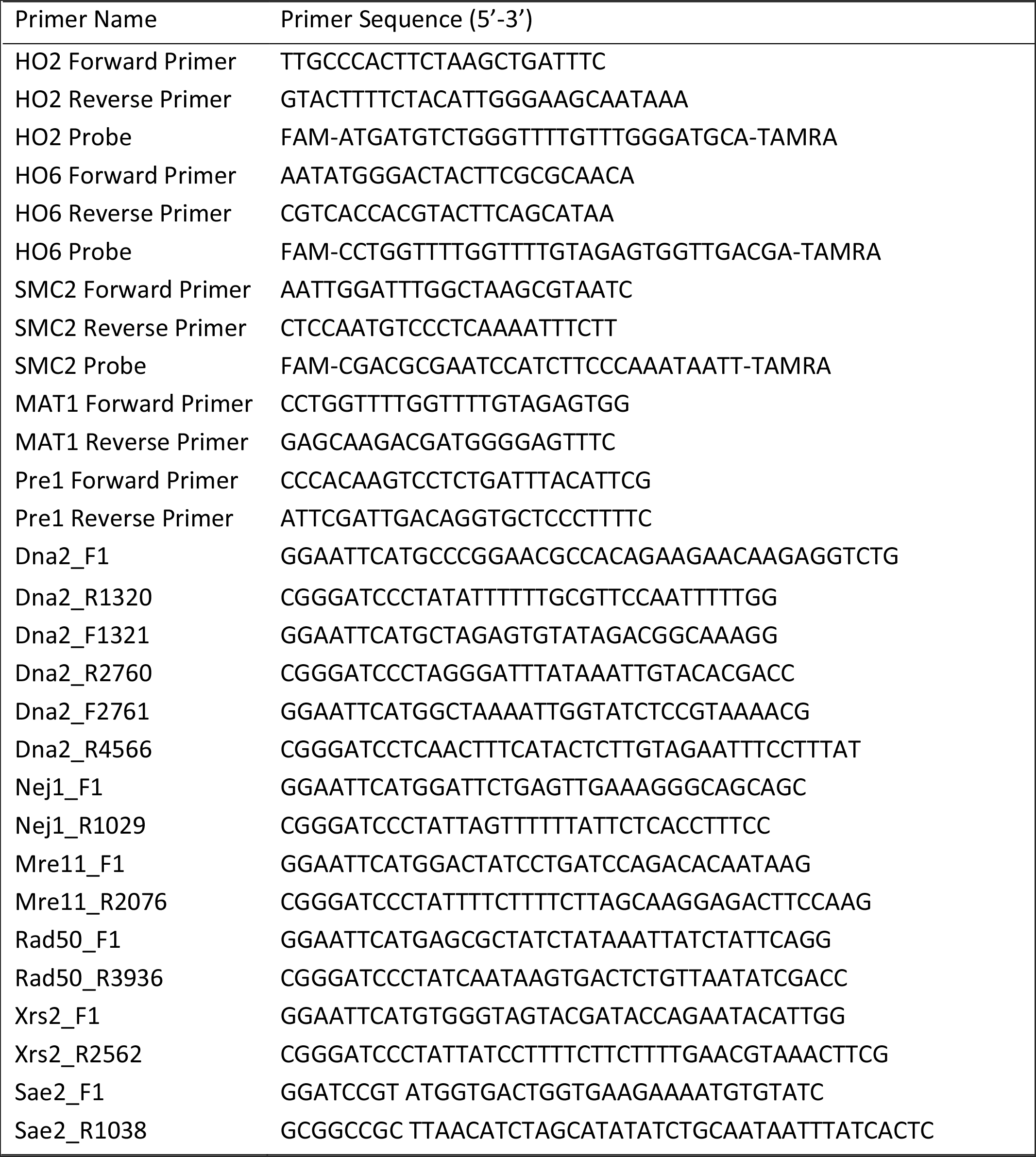
Primers and Probes used in this study.

